# Human Vascularized Macrophage-Islet Organoids to Model Immune-Mediated Pancreatic β cell Pyroptosis upon Viral Infection

**DOI:** 10.1101/2024.08.05.606734

**Authors:** Liuliu Yang, Yuling Han, Tuo Zhang, Xue Dong, Jian Ge, Aadita Roy, Jiajun Zhu, Tiankun Lu, J. Jeya Vandana, Neranjan de Silva, Catherine C. Robertson, Jenny Z Xiang, Chendong Pan, Yanjie Sun, Jianwen Que, Todd Evans, Chengyang Liu, Wei Wang, Ali Naji, Stephen C.J. Parker, Robert E. Schwartz, Shuibing Chen

## Abstract

There is a paucity of human models to study immune-mediated host damage. Here, we utilized the GeoMx spatial multi-omics platform to analyze immune cell changes in COVID-19 pancreatic autopsy samples, revealing an accumulation of proinflammatory macrophages. Single cell RNA-seq analysis of human islets exposed to SARS-CoV-2 or Coxsackievirus B4 (CVB4) viruses identified activation of proinflammatory macrophages and β cell pyroptosis. To distinguish viral versus proinflammatory macrophage-mediated β cell pyroptosis, we developed human pluripotent stem cell (hPSC)-derived vascularized macrophage-islet (VMI) organoids. VMI organoids exhibited enhanced marker expression and function in both β cells and endothelial cells compared to separately cultured cells. Notably, proinflammatory macrophages within VMI organoids induced β cell pyroptosis. Mechanistic investigations highlighted TNFSF12-TNFRSF12A involvement in proinflammatory macrophage-mediated β cell pyroptosis. This study established hPSC-derived VMI organoids as a valuable tool for studying immune cell-mediated host damage and uncovered mechanism of β cell damage during viral exposure.

## INTRODUCTION

A strong connection between Coronavirus disease 19 (COVID-19) and diabetes is now recognized. Since the beginning of the pandemic, there have been reports of new-onset diabetes^1–5^ and exacerbated complications in patients with pre-existing diabetes. Moreover, a rise in type 1 diabetes (T1D) incidence has been observed^4,6^. A study from the Centers for Disease Control and Prevention reported that persons aged <18 years with COVID-19 were more inclined to receive a new diabetes diagnosis compared to those without COVID-19. Studies reported a heightened T1D and type 2 diabetes (T2D) incidence rates after the beginning of pandemic, surpassing pre-pandemic period^7,8,9^. In addition to SARS-CoV-2, a number of studies suggests the correlation between viral infections and T1D^10^, including enteroviruses^11^, such as coxsackievirus B^12,13^, as well as rotavirus^14^, mumps virus^15^, and cytomegalovirus^16^. Coxsackievirus B4 (CVB4), a positive-sense single-stranded RNA virus, isolated from newly diagnosed T1D patients could infect and induce destruction of human islet cells *in vitro*^17^.

In infectious diseases, multiple mechanisms contribute to the observed host injury. Our group and others discovered that SARS-CoV-2 infection induces the transdifferentiation of human β cells^18^ and damage of β cells^19,20^. In addition, the accumulation of macrophages has been reported in the lungs^21^ and heart^22^ of COVID-19 patients. Further insight would benefit from robust human models to explore immune cell-mediated host damage. Human pluripotent stem cells (hPSCs)^23^ provide a powerful *in vitro* platform for studying disease mechanisms, developing cell therapy approaches and drug screening^24–26^. Many efforts have applied hPSC-based platforms to study SARS-CoV-2 tropism^27^ and host responses. Recently, we performed a 2D co-culture system utilizing hPSC-derived cardiomyocytes and macrophages and identified a Janus kinase (JAK) inhibitor that effectively thwarts macrophage-mediated damage to cardiac cells^22^.

In this study, we applied spatial multi-omics assays to comprehensively analyze pancreatic autopsy samples of COVID-19 patients and identified the accumulation of proinflammatory macrophages in COVID-19 samples. Single cell RNA-seq analysis confirmed the activation of proinflammatory macrophages and enrichment of the pyroptotic pathway in β cells of human islets exposed to SARS-CoV-2 or CVB4 viruses. Next, we developed a vascularized macrophage-islet (VMI) organoid model containing hPSC-derived endocrine cells, macrophages, and endothelial cells and found that proinflammatory macrophages induced β cell pyroptosis through the secretion of IL-1β and interaction with β cells via the TNFSF12-TNFRSF12A pathway. This study not only establishes a VMI organoid model to study macrophage-mediated host damage, but also identifies the previously unknown role of TNFSF12-TNFRSF12A-mediated pyroptosis in β cell damage in infectious diseases.

## RESULTS

### Spatial multi-omics analysis to identify the activation of proinflammatory macrophages in pancreatic autopsy samples of COVID-19 patients

To systematically analyze the pancreatic damage of COVID-19 patients, we collected pancreatic autopsy samples from 7 COVID-19 patients and 8 age and gender-matched control subjects (**Table S1**). GeoMx multi-omics assays were applied to analyze two adjacent tissue sections of each donor, providing paired analysis of changes at both transcriptome and protein levels (**Figure S1A**). For GeoMx analysis, we selected 6 regions of interests (ROIs) in the islet area, 3 ROIs in ductal area and 3 ROIs in exocrine area per sample (**Figure 1A and Figure S1A**). Through morphology marker staining (INS/Pan-CK/TOTO-3), we observed no obvious change of islet areas, but lower percentage of INS^+^ cells within islets of COVID-19 samples compared to control samples (**Figures 1B and 1C**), which is consistent with the previous reports^18^. 3D PCA analysis of whole transcriptome sequencing data from ROIs in the islet areas showed distinct transcriptional profiles in COVID-19 samples separated from control samples (**Figure 1D**). Pathway analysis of differentially expressed (DE) genes between ROIs in islet areas from COVID-19 and control samples highlighted viral infection associated pathways, such as viral mRNA translation, influenza infection, interferon α/β signaling pathways and stress associated pathways, such as cellular response to stress or external stimuli pathways, and toll-like receptor 2 cascade pathway (**Figure 1E**). Consistently, PCA of ROIs in ductal and exocrine areas also showed separation between COVID-19 and control samples (**Figures S1B and S1C**). Moreover, pathway analysis of DE genes between ROIs in ductal and exocrine areas in COVID-19 and control samples also revealed the enrichment of interferon signaling pathways in COVID-19 samples (**Figures S1D and S1E**).

**Figure 1.**
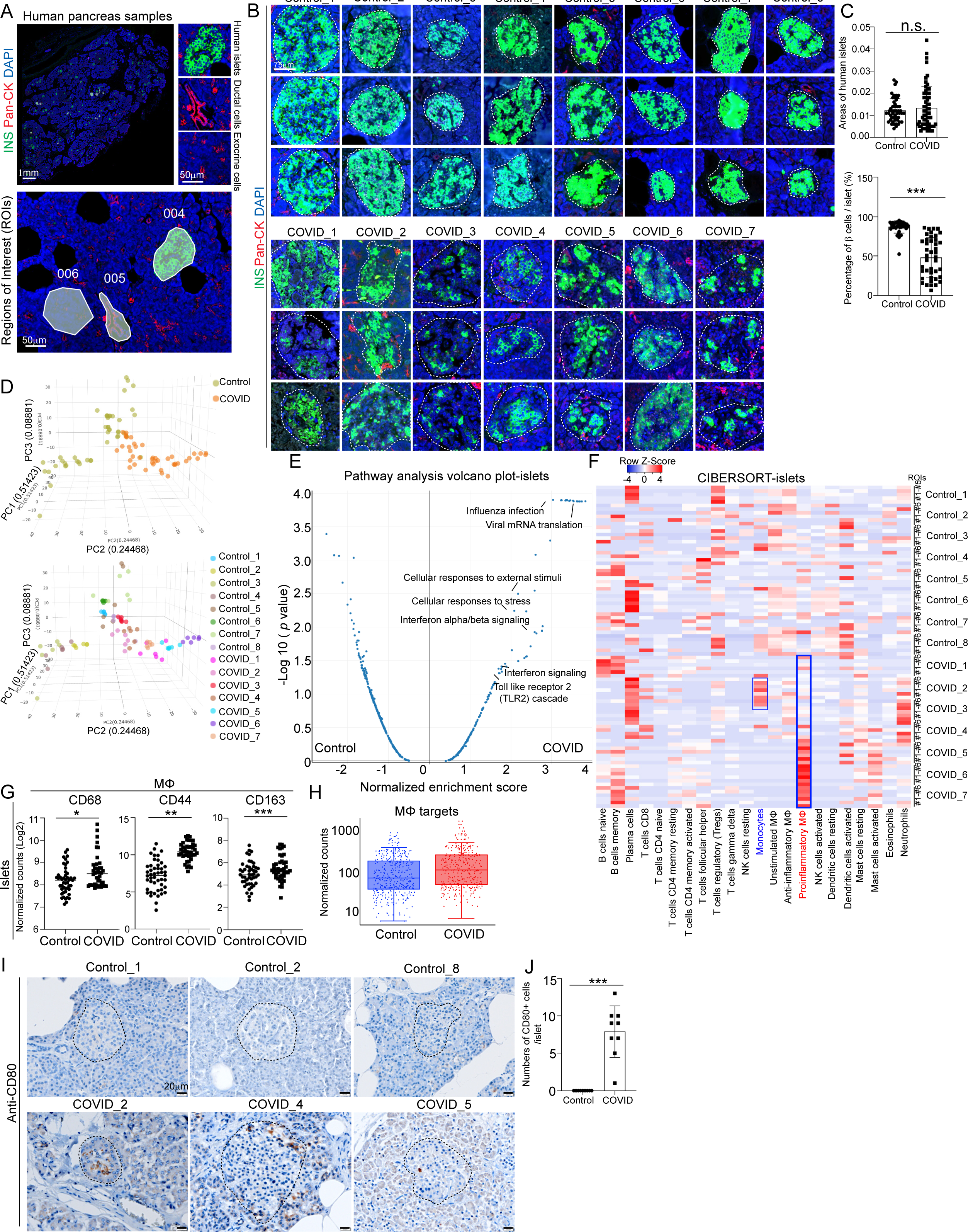
Macrophage accumulation in islets of COVID-19 pancreatic autopsy samples. **(A)** Representative images illustrating morphology marker and selection of ROIs using GeoMx platform. 004: islet area; 005: ductal area; 006: exocrine area. Scale bar= 1 mm or 50 µm. **(B)** Representative images illustrating the insulin (INS) staining in COVID-19 (N=7) and control (N=8) pancreatic autopsy samples. Dotted lines encircle the islet regions. Scale bar=75 µm. **(C)** Quantification of areas of islets and percentages of INS^+^ β cells per islet in COVID-19 (N=7) and control (N=8) pancreatic autopsy samples. **(D)** 3D PCA plot of data from human islet areas of COVID-19 (N=7) and control (N=8) pancreatic autopsy samples. **(E)** Volcano plot of transcriptome sequencing data highlighting the pathways enriched in human islet areas of COVID-19 (N=7) versus control (N=8) pancreatic autopsy samples. **(F)** Heatmap of the CIBERSORT analysis of immune cells (LM22) using the GeoMx whole transcriptome sequencing data of human islet areas of COVID-19 (N=7) and control (N=8) pancreatic autopsy samples. **(G)** Normalized counts (Log2) of marker proteins associated with macrophages from human islet areas of COVID-19 (N=7) and control (N=8) pancreatic autopsy samples. Each dot represents one count in each ROI. **(H)** Box plot of normalized counts of macrophage associated targets in human islet areas of COVID-19 (N=7) and control (N=8) pancreatic autopsy samples. Each dot represents one count in each ROI. **(I and J)** Immunohistochemistry staining (I) and quantification (J) of CD80 in COVID-19 (N=3) and control (N=3) pancreatic autopsy samples. Dotted lines encircle the regions of the islets. Scale bar=20 μm. *P* values were calculated by unpaired two-tailed Student’s t test. n.s., no significance; \**P* < 0.05, \*\**P* < 0.01, \*\*\**P* < 0.001. See also Figure S1.

To further analyze the changes in immune cell composition, we conducted CIBERSORT analysis of transcriptome profiles from COVID-19 and control samples. Within the ROIs in islet area of the 7 COVID-19 samples, we found 4 samples (#4-#7) enriched with proinflammatory macrophages, and 2 samples (#2-#3) enriched with monocytes (**Figure 1F**). The enrichment of monocytes or proinflammatory macrophages were not dependent on the pre-existing type 2 diabetes condition of the subjects (**Figure S1F**). Consistently, we detected enrichment of proinflammatory macrophages in ductal ROIs in 4 (#4-#7) out of 7 COVID-19 samples, and monocyte enrichment in ductal ROIs in 1 (#3) out of 7 COVID-19 samples (**Figure S1G**). In exocrine ROIs, we observed enrichment of proinflammatory macrophages in 4 (#4-#7) out of 7 COVID-19 samples (**Figure S1H**).

We further conducted GeoMx protein assays and found that macrophages were enriched in islet ROIs of COVID-19 samples compared to control samples (**Figure 1G and 1H**); while T cells, NK cells, B cells and neutrophils were not enriched (**Figure S1I**). Moreover, the proteins related to T cell activation were not increased in islet ROIs of COVID-19 samples compared to control samples (**Figure S1J**). Notably, CD44, previously reported to regulate the TLR2-mediated macrophage activation and proinflammatory responses^28,29^, was also found to be significantly increased in ROIs in islet, ductal, and exocrine areas of COVID-19 samples (**Figure 1G and Figure S1K**). CD163, which functions as the scavenger receptor is highly upregulated in infiltrating macrophages in sites of inflammation^30,31^. Soluble CD163, was also identified as a biomarker of macrophage activation and associated with type 2 diabetes mellitus (T2DM), insulin resistance, and β cell dysfunction^32^. Finally, immunohistochemistry validated the accumulation of both CD163^+^ macrophages and CD80^+^ proinflammatory macrophages in pancreatic tissues of COVID-19 patients (**Figures 1I-1J and Figures S1L-S1M**).

### Single cell RNA-seq analysis identifies activation of proinflammatory macrophages and β cell pyroptosis in SARS-CoV-2 or CVB4 exposed human islets

To further explore the status of macrophages upon virus exposure in human islets, we performed single cell RNA-seq (scRNA-seq) of human islets upon exposure of SARS-CoV-2 or CVB4. UMAP analysis revealed nine cell clusters within human islets (**Figures S2A and S2B**). In our previous publication, we have already characterized the SARS-CoV-2 infected human islets^33^. Here, we further characterized the CVB4 infected human islets. UMAP and violin plot showed the high expression of CVB4 virus *polyprotein* in endocrine cells (β cells, α cells and δ cells), as well as mesenchymal cells, immune cells, and endothelial cells (**Figures S2C and S2D**). Immunostaining confirmed the colocalization of enterovirus (CVB4) and endocrine cell markers, including INS (β cells), GCG (α cells) and SST (δ cells) (**Figures S2E-S2I**).

We then focused on the immune cell population and performed sub-clustering analysis, identifying five sub-clusters (**Figure 2A**). UMAP and violin plots confirmed the expression of marker genes for each subpopulation (**Figure 2B**). We compared the transcriptional profiles of macrophages and found increased expression of proinflammatory macrophage-associated genes, including *IL1B*, *IL6*, *CXCL8* and *TNF* in macrophages of human islets upon SARS-CoV-2 exposure (**Figure 2C**). Immunostaining also confirmed the activation of proinflammatory macrophages in human islets upon SARS-CoV-2 infection (**Figures 2D-2E and Figures S2J-S2M**). We further analyzed several cell death-associated pathways within β cell cluster of human islets exposed to SARS-CoV-2 virus. Interestingly, we found the activation of pyroptosis and apoptosis pathways in β cells of human islets exposed to SARS-CoV-2 (**Figure 2F**). Our previous studies have reported the activation of apoptosis of SARS-CoV-2 infected β cells^27^. In the current study, we focused on β cell pyroptosis. Dot plot analysis showed increased expression levels of pyroptosis-associated genes in both SARS-CoV-2+ and SARS-CoV-2- β cells of human islets exposed to SARS-CoV-2 (**Figure 2G and Figure S3A**). Immunostaining further confirmed the increased expression of cleaved caspase1 (CASP1) in β cells of human islets upon SARS-CoV-2 infection (**Figures 2H and 2I**). In addition, we found enrichment of pyroptosis pathway in other endocrine cell clusters (α and δ cell clusters) of human islets exposed to SARS-CoV-2 (**Figure S3B**). Moreover, autophagy pathway was enriched in mesenchymal cell cluster; while ferroptosis and apoptosis pathways were enriched in endothelial cell cluster of human islets exposed to SARS-CoV-2 (**Figure S3B**).

**Figure 2.**
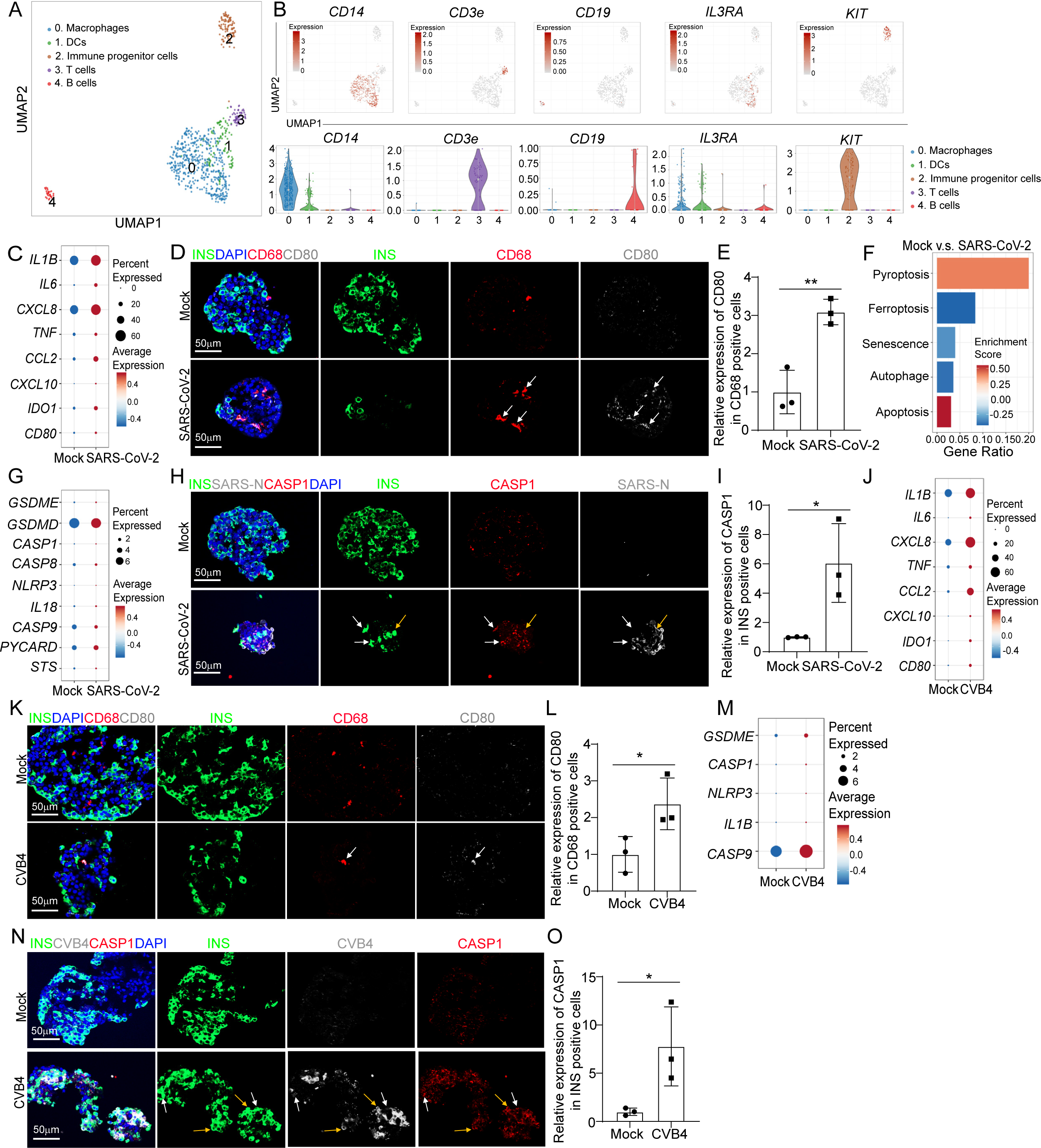
Single cell RNA-seq analysis of human islets upon SARS-CoV-2 or CVB4 exposure. **(A)** UMAP of immune cell populations in human islets exposed to mock, SARS-CoV-2 (MOI=1) or CVB4 (2X10^6^ PFU/ml) . **(B)** UMAP and violin plots of immune cell markers. **(C)** Dot plot analysis of proinflammatory macrophage-associated genes in macrophages of human islets exposed to mock or SARS-CoV-2 (MOI=1). **(D and E)** Confocal images (D) and quantification (E) of the relative expression of CD80 in CD68^+^ cells in human islets exposed to mock or SARS-CoV-2 (MOI=0.5). The white arrows highlight the CD68^+^CD80^+^ cells. Scale bar= 50 µm. **(F)** Pathway enrichment analysis of cell death pathways in β cell cluster of human islets exposed to mock or SARS-CoV-2 (MOI=1). **(G)** Dot plot analysis of pyroptosis associated genes in β cell cluster of human islets exposed to mock or SARS-CoV-2 (MOI=1). **(H and I)** Confocal images (H) and quantification (I) of the relative expression of CAPS1 in human islets exposed to mock or SARS-CoV-2 (MOI=0.5). The yellow arrows highlight the expression of CASP1 in SARS-N^+^INS^+^ cells while the white arrows highlight the expression of CASP1 in SARS-N^-^INS^+^ cells. Scale bar= 50 µm. **(J)** Dot plot analysis of proinflammatory macrophage-associated genes in macrophage cluster of human islets exposed to mock or CVB4 (2x10^6^ PFU/ml). **(K and L)** Confocal images (K) and quantification (L) of the relative expression of CD80 in CD68^+^ cells in human islets exposed mock or CVB4 (2x10^6^ PFU/ml). The white arrows highlight the CD68^+^CD80^+^ cells. Scale bar= 50 µm. **(M)** Dot plot analysis of pyroptosis pathway associated genes in β cell cluster of human islets exposed to mock or CVB4 (2x10^6^ PFU/ml). **(N and O)** Fluorescent images (N) and quantification (O) of the relative expression of CAPS1 in human islets exposed to mock or CVB4 (2x10^6^ PFU/ml). The yellow arrows highlight the expression of CASP1 in SARS-N^+^INS^+^ cells while the white arrows highlight the expression of CASP1 in SARS-N^-^INS^+^ cells. Scale bar= 50 µm. N=3 independent biological replicates. Data was presented as mean ± STDEV. *P* values were calculated by unpaired two-tailed Student’s t test. \**P* < 0.05, \*\**P* < 0.01. See also Figure S2 and S3.

Next, we analyzed human islets exposed to CVB4 virus. Similar with SARS-CoV-2 virus, CVB4 virus exposure also induced activation of proinflammatory macrophages (**Figures 2J-2L and Figures S3C-S3F**). Furthermore, dot plots showed increased expression of pyroptotic pathway associated genes in both CVB4+ and CVB4- β cells of human islets upon CVB4 infection (**Figure 2M and Figure S3G**). Finally, immunostaining confirmed the increased expression of CASP1 in β cells of human islets upon CVB4 infection (**Figures 2N and 2O**). Together, these data demonstrate the activation of proinflammatory macrophages and β cell pyroptosis in human islets exposed to SARS-CoV-2 or CVB4 viruses.

To further analyze the changes of immunogenicity profiles of β cells in response to viral infection, we analyzed the expression of HLA molecules and autoantigen associated genes. We observed a pattern of the increased expression of HLA class I genes in β cells of human islets exposed to SARS-CoV-2 versus mock (**Figure S3H**). In contrast, there was a trend of reduced expression of HLA class I genes in β cells of human islets exposed to CVB4 versus mock (**Figure S3I**). In terms of autoantigen expression, we observed a similar increase in the expression of *GAD2* and *IAPP* in β cells of human islets exposed to SARS-CoV-2 or CVB4 compared to mock conditions. For *CHGA* and *SLC30A8*, we noted different trends of expression, which increased in β cells of human islets exposed to SARS-CoV-2 but decreased in β cells of human islets exposed to CVB4 (**Figures S3J and S3K**). Moreover, we also examined the genes related to antigen presentation and found an increased expression of antigen presentation associated genes in β cells of human islets exposed to SARS-CoV-2 and a reduced expression of them in β cells of human islets exposed to CVB4 (**Figures S3L and S3M**).

### Construction of a vascularized macrophage-islet organoid

To determine whether β cell pyroptosis is caused by proinflammatory macrophages activation, we constructed a vascularized macrophage-islet organoid (VMI organoid) model (**Figure 3A**). First, we differentiated MEL-1*^INS/GFP^* hESCs into pancreatic endocrine cells (**Figure S4A**). At day 16, we detected the robust generation of INS^+^ β cells, GCG^+^ α cells and SST^+^ δ cells (**Figure S4B**). H9 hESCs were differentiated toward macrophages which expressed CD11B, CD14 and CD206, but not CD80 (**Figures S4C- S4D**). Functional assays confirmed that hESC-derived macrophages can engulf bacteria, indicating that they exhibited phagocytic activity which similar to primary human macrophages (**Figure S4E**). Human islets are highly vascularized and endothelial cells play an important role in systemic inflammatory responses^34,35^, as well as pancreatic cell development^36–38^. Thus, we decided to add endothelial cells to VMI organoids. ETV2 was reported to promote the development of endothelial cells^39,40^. Here, we overexpressed ETV2 to promote the differentiation and function of endothelial cells from H1 hESCs^40–42^ (**Figure S4F**). qRT-PCR confirmed the overexpression of ETV2 in H1 hESCs (**Figure S4G**) and immunostaining confirmed that differentiated endothelial cells expressed PECAM1 (CD31) (**Figure S4H**).

**Figure 3.**
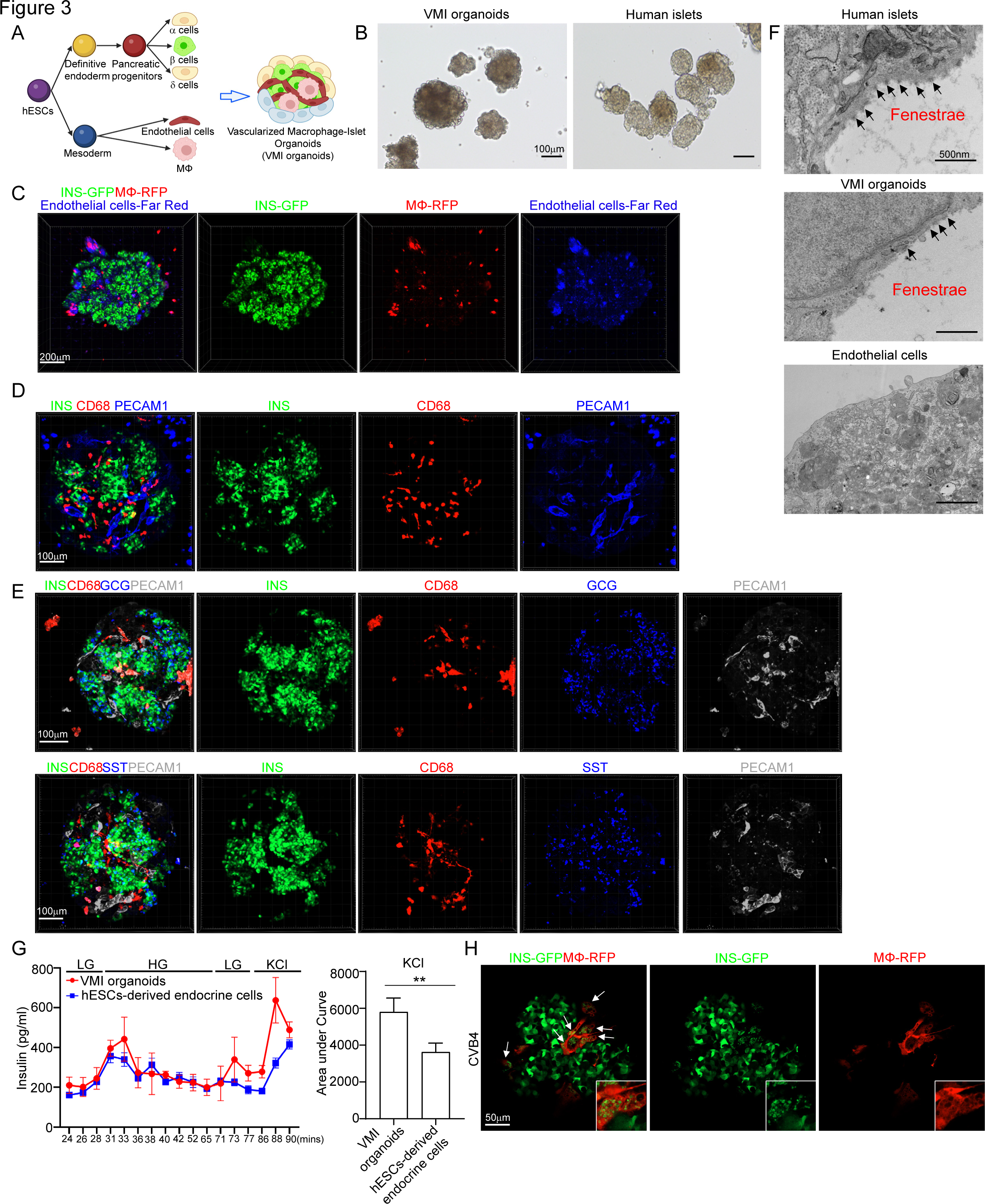
Construction of hPSC-derived VMI organoids. **(A)** Schematic representation of VMI organoids construction. **(B)** Phase contract images of VMI organoids at day 14 after reaggregation and human islets. Scale bar= 200 µm. **(C)** Composite Z-stack confocal images of live VMI organoids at day 14 after reaggregation. β cells: INS-GFP; macrophages: RFP; and endothelial cells: Far Red. Scale bar= 200 µm. **(D)** Composite Z-stack confocal images of VMI organoids at day 14 after reaggregation stained with antibodies against INS, CD68 and PECAM1 (CD31). Scale bar= 100 µm. **(E)** Composite Z-stack confocal images of VMI organoids at day 14 after reaggregation stained with antibodies against INS, CD68, GCG, SST and PECAM1 (CD31). Scale bar= 100 µm. **(F)** Transmission electron microscope (TEM) images of human islets, VMI organoids at day 14 after reaggregation, and endothelial cells without reaggregation. Arrows indicate fenestrae. Scale bar= 500 nm. **(G)** Dynamic glucose stimulated insulin secretion of VMI organoids at day 14 after reaggregation and hPSC-derived endocrine cells. LG (low glucose): 2 mM D-glucose; HG (high glucose): 20 mM D-glucose; KCl: 30 mM KCl. Quantification was performed using the areas under curve of KCl stimulation from 86 min to 90 min. **(H)** Composite Z-stack confocal images of VMI organoids at day 7 after reaggregation upon CVB4 infection (2x10^6^PFU/ml). β cells: INS-GFP, macrophages: RFP. Scale bar= 50 µm. Arrows highlight RFP^+^ macrophages that have phagocytosed damaged INS-GFP^+^ β cells. N=3 independent biological replicates. Data was presented as mean ± STDEV. *P* values were calculated by unpaired two-tailed Student’s t test. \*\**P* < 0.01. See also Figure S4.

After optimizing the culture medium and cell ratio, we mixed the hESC-derived endocrine cells, unstimulated macrophages, and endothelial cells in three-dimension (3D) culture to form organoids (**Figure 3A**). The VMI organoids exhibited similar size and morphologies as primary human islets (**Figure 3B**). To perform live imaging of the VMI organoid, we labeled cells with fluorescent reporters or CellTrace dye. Pancreatic endocrine cells were derived from MEL-1*^INS/GFP^*hESCs, allowing real-time monitoring of INS-GFP^+^ β cells.

Macrophages were derived from RFP labelled H9 (RFP-H9) hESCs and purified using magnetic sorting before organoid formation. Additionally, H1 hESC-derived endothelial cells were selected by magnetic sorting and the purified endothelial cells were labelled with CellTrace proliferative far-red dye before forming 3D organoids. 3D confocal images confirmed the presence of INS-GFP^+^ β cells, RFP^+^ macrophages, and far red^+^ endothelial cells in VMI organoids (**Figure 3C and Supplemental Video 1**). Immunostaining of VMI organoids confirmed the presence of INS^+^ β cells, CD68^+^ macrophages and PECAM1^+^ endothelial cells (**Figure 3D and Supplemental Video 2**). Most of the INS^+^ β cells in VMI organoids co-expressed NKX6.1, a key transcription factor of β cells (**Figure S4I and Supplemental Video 3**). Immunostaining further confirmed the presence of GCG^+^ α cells and SST^+^ δ cells in VMI organoids (**Figure 3E and Supplemental Videos 4-5**).

Next, we used different assays to determine whether the cells in VMI organoids closely resembled the cells in primary islets. Initially, electron microscopy (EM) was used to observe fenestrae, which are transcellular pores found in endothelial cells facilitating the transfer of substances between blood and the extravascular space^43^. Indeed, the fenestrations were detected in the endothelial cells of both primary human islets and VMI organoids, but not in separately cultured endothelial cells (**Figure 3F**). We performed an acetylated-low density lipoprotein (Ac-LDL) uptake assay to assess the function of endothelial cells in VMI organoids. Ac-LDL can bind to the receptor on the surface of vascular endothelial cells, facilitating the delivery of cholesterol via endocytosis^44,45^. We found co-localization of Ac-LDL with PECAM1^+^ endothelial cells (**Figure S4J**). Then, we performed dynamic glucose-stimulated insulin secretion (GSIS) to examine the secretion of insulin upon glucose or KCl stimulation. We found increased insulin expression in VMI organoids than separately cultured hESC-derived endocrine cells under both high glucose and KCl stimulation conditions. The amount of insulin secreted upon KCl stimulation was significantly higher in VMI organoids than separately cultured hESC-derived endocrine cells (**Figure 3G**). Besides, we also found a decrease of GCG secretion in VMI organoids compared to separately cultured endocrine cells (**Figure S4K**). To further elevate the function of β cells upon low glucose and high glucose stimulation, we performed dynamic calcium Flu4 imaging. We detected dynamic calcium mobilization in cells of VMI organoids upon high glucose stimulation (**Figure S4L and Supplemental Video 6**). Together, the data indicate that the pancreatic β cells and endothelial cells in VMI organoids are functionally more mature than cells that are cultured separately.

Finally, we exposed the VMI organoids with CVB4 virus and found macrophages engulfing the damaged β cells upon virus infection (**Figure 3H and Supplemental Video 7**). To examine monocyte infiltration, we created organoids containing endocrine cells and endothelial cells (VI organoids) and monitored monocyte infiltration upon CVB4 infection. We first added monocytes to VI organoids, then introduced CVB4, and conducted live cell imaging at 24hpi and 48hpi. We did not find obviously infiltration of monocytes into VI organoids (**Figure S4M**).

### Single cell multi-omics analysis of VMI organoids

We then performed scRNA-seq and single nucleus assay for transposase-accessible chromatin using sequencing (snATAC-seq) to compare the cell compositions, transcriptional and epigenetic profiles of VMI organoids and separately cultured cells^46^ (**Figure 4A**). Cells that were cultured in separate plates but mixed together before library preparation was compared to cells in VMI organoids (**Figure S4N**). UMAP analysis identified 9 cell clusters (**Figure 4A**). Dot plot of scRNA-seq analysis (**Figure 4B**) and integrative genomics viewer plot of snATAC-seq (**Figure S4O**) confirmed the marker gene expression in each cluster. Consistent with previous studies^47–49^, *GCG* expression was detected in β cell cluster and *INS* expression was detected in α and δ cell clusters, suggesting the immature status of hESC-derived endocrine cells. Next, we compared the cells in VMI organoids with separately cultured cells (**Figure 4C**). Pie chart showed the relative proportions of major cell types in VMI organoids (**Figure 4D**). Volcano plot analysis of gene expression in β cell cluster showed decreased expression of non-β cell associated genes, *AFP*^50^, *GCG*, *SST*, *ACTB* and *PRSS2* and increased expression of β cell associated genes *RPL13A*^51^ and *SMIM32*^52^ in β cell cluster of VMI organoids compared to separately cultured cells (**Figure 4E**). Dot plot and violin plot analysis also showed that genes associated with β cell identity and function, including *SLC2A1*^53–55^, *PIK3CB*^56^, *HNF1B*, *PAX6*, *PDX1* and *INS*, are relatively increased in β cell cluster of VMI organoids (**Figure 4F and Figure S4P**). Consistently, snATAC-seq analysis showed increased open chromatin accessibility peaks of *SLC2A1*^53–55^, *INS* and *PDX1* in β cells of VMI organoids compared to separately cultured cells which might indicate the potential of increased gene expression of *SLC2A1*, *INS* and *PDX1* (**Figure 4G**). In addition, dot plot and violin plot also revealed the upregulation of genes associated with endothelial cell function in endothelial cell cluster of VMI organoids compared to separately cultured cells, including *INSR*^43^, *VWF*^57^, *PDGFB*^58^, *EDN1*^59^, *S1PR1*^60^ and *RSPO3*^61^ (**Figure 4H and S4Q**).

**Figure 4.**
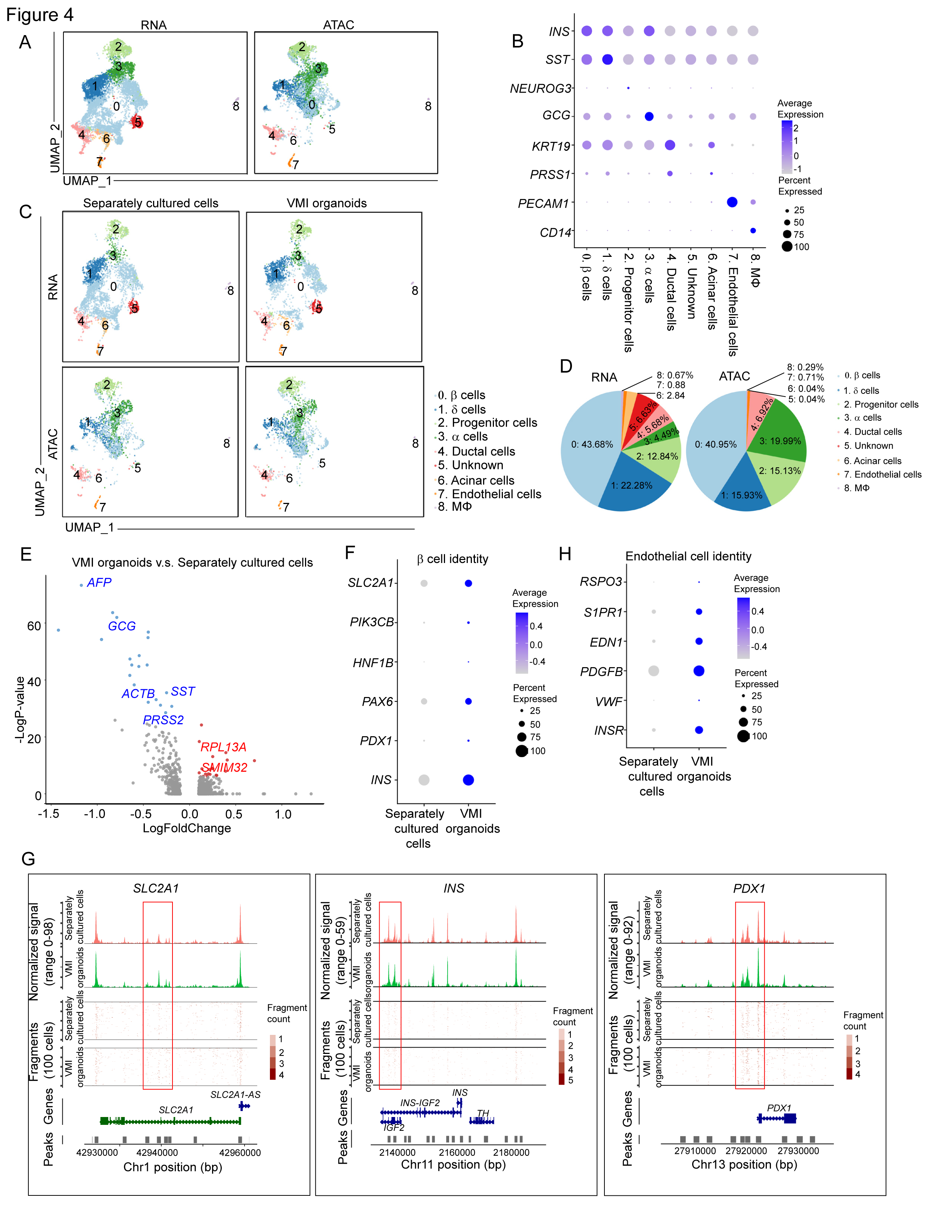
Single cell multi-omics analysis of VMI organoids. (A) Integrative UMAP of scRNA-seq and snATAC-seq analysis of VMI organoids at day 7 after reaggregation and separately cultured cells. (B) Dot plot displaying cell markers of each cluster using scRNA-seq dataset. (C) Individual UMAP of scRNA-seq and snATAC-seq analysis of VMI organoids at day 7 after reaggregation and separately cultured cells. (D) Pie chart showed the relative percentages of each cell types in VMI organoids at day 7 after reaggregation. (E) Volcano plot of DE genes in β cell cluster of VMI organoids at day 7 after reaggregation versus separately cultured cells. (F) Dot plot analysis of β cell associated genes in β cell cluster of VMI organoids at day 7 after reaggregation and separately cultured cells. (G) Chromatin accessibility signals of *SLC2A1*, *INS*, *PDX1* in β cell cluster of VMI organoids at day 7 after reaggregation and separately cultured cells. The normalized signal shows the averaged frequency of sequenced DNA fragments within a genomic region. The fragment shows the frequency of sequenced fragments within a genomic region for individual cells. (H) Dot plot analysis of endothelial cell associated genes in endothelial cell cluster of VMI organoids at day 7 after reaggregation and separately cultured cells. See also Figure S4.

### Proinflammatory macrophages cause β cell pyroptosis

We have shown the activation of proinflammatory macrophages, as well as upregulation of the pyroptotic pathway in β cell cluster of human islets exposed to SARS-CoV-2 or CVB4 viruses. Here, we also detected the activation of proinflammatory macrophages, as well as β cells pyroptosis, in VMI organoids exposed to SARS-CoV-2 or CVB4 viruses (**Figures S5A-S5D**). To determine whether proinflammatory macrophages cause β cell pyroptosis, we constructed the VMI organoids with proinflammatory or unstimulated macrophages. First, LPS and IFN-γ were used to stimulate macrophages into proinflammatory status (**Figure S5E**). Both RNA-seq and ELISA analysis confirmed the increased expression of proinflammatory associated genes and cytokines, including IL- 1β and IL-6 in proinflammatory macrophages (**Figures S5F and S5G**). Then, we constructed VMI organoids using either unstimulated or proinflammatory macrophages. VMI organoids containing proinflammatory macrophages showed decreased expression levels of INS compared to VMI organoids containing unstimulated macrophages (**Figures 5A-5B and Supplemental Video 8, 9**). We collected the supernatant of VMI organoids containing proinflammatory or unstimulated macrophages and confirmed the increased expression of IL-1β, IL-6 and TNF-α in the supernatant of VMI organoids containing proinflammatory macrophages (**Figure 5C**).

**Figure 5.**
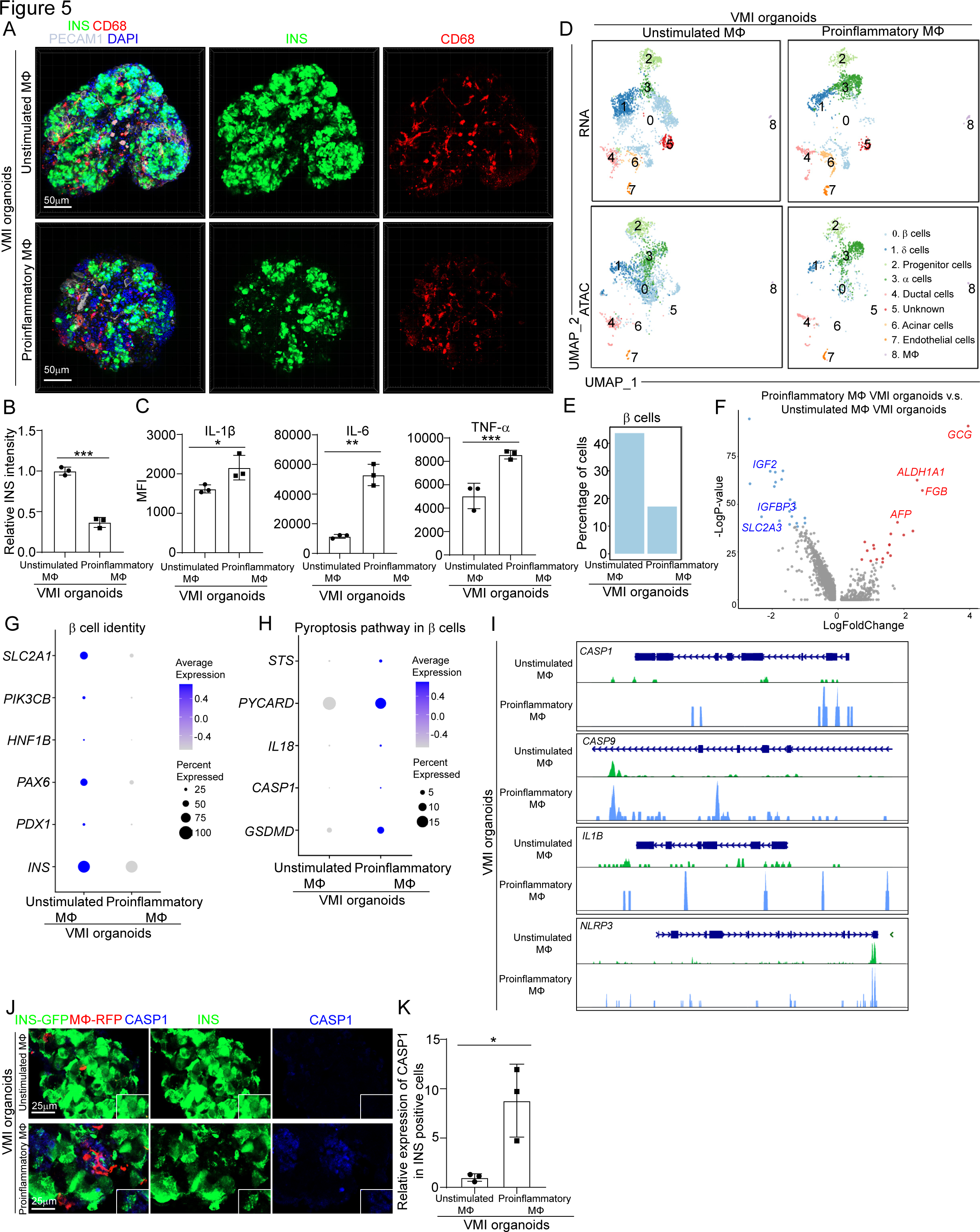
Construction and multi-omics analysis of VMI organoids containing unstimulated and proinflammatory macrophages. **(A and B)** Composite Z-stack confocal images (A) and quantification (B) of INS intensity in INS^+^ cells of VMI organoids at day 7 after reaggregation containing unstimulated or proinflammatory macrophages stained with the antibodies against INS, CD68 and PECAM1 (CD31). Scale bar= 50 µm. (C) Measurements of cytokine secretions in the supernatant of VMI organoids at day 5 after reaggregation containing unstimulated or proinflammatory macrophages. (D) Integrative UMAP of VMI organoids at day 7 after reaggregation containing unstimulated or proinflammatory macrophages. (E) Percentage of cells in β cell cluster in VMI organoids at day 7 after reaggregation containing unstimulated or proinflammatory macrophages. (F) Volcano plot of DE genes in β cell cluster of VMI organoids at day 7 after reaggregation containing proinflammatory versus unstimulated macrophages. (G) Dot plot analysis of β cell identity associated genes in β cell cluster of VMI organoids at day 7 after reaggregation containing unstimulated or proinflammatory macrophages. (H) Dot plot analysis of pyroptosis pathway associated genes in β cell cluster of VMI organoids at day 7 after reaggregation containing unstimulated or proinflammatory macrophages. (I) Chromatin accessibility signals of *CASP1*, *CASP9, IL1B* and *NLRP3* in β cell cluster of VMI organoids at day 7 after reaggregation containing unstimulated or proinflammatory macrophages. The normalized signal shows the averaged frequency of sequenced DNA fragments within a genomic region. The fragment shows the frequency of sequenced fragments within a genomic region for individual cells. **(J and K)** Immunostaining (J) and quantification (K) of CASP1 staining in INS+ cells of VMI organoids at day 7 after reaggregation containing unstimulated or proinflammatory macrophages. β cells: INS-GFP, macrophages: RFP, endothelial cells: Far Red. CASP1: grey. Scale bar= 25 µm. N=3 independent biological replicates. Data was presented as mean ± STDEV. *P* values were calculated by unpaired two-tailed Student’s t test. \**P* < 0.01, \*\**P* < 0.05, \*\*\**P* < 0.001. See also Figure S5.

Next, scRNA-seq and snATAC-seq were performed to analyze the VMI organoids containing proinflammatory or unstimulated macrophages. Consistent with the previous analysis, 9 cell clusters were identified in VMI organoids. UMAP showed a decrease of the β cell cluster in VMI organoids with proinflammatory macrophages, which was also confirmed by quantification of the percentage of β cells (**Figures 5D and 5E**). Moreover, volcano plot comparing the β cell cluster of VMI organoids containing proinflammatory macrophages to that of organoids containing unstimulated macrophages showed the downregulation of β cell identity and function associated genes, including the decreased expression levels of *IGF2*^62^, *IGFBP3*^63^ and *SLC2A3*^64^, and upregulation of non-β cell identity and function associated genes, *GCG*, *ALDH1A1*^65^*, FGB*^66^ and *AFP*^50^ (**Figure. 5F**). Dot plot analysis further confirmed the downregulation of β cell identity and function associated genes, including *SLC2A1*^53–55^, *PIK3CB*,^56^ *HNF1B*, *PAX6*, *PDX1* and *INS,* in the β cluster of VMI organoids containing proinflammatory macrophages compared to VMI organoids containing unstimulated macrophages (**Figure 5G**). Consistently, snATAC-seq analysis also showed decreased open chromatin accessibility peaks of *INS* and *PDX1* in β cells of VMI organoids containing proinflammatory macrophages (**Figure S5H**). Furthermore, dot plot analysis of scRNA-seq analysis also showed increased expression of pyroptotic pathway associated genes in the β cluster of VMI organoids containing proinflammatory macrophages (**Figure 5H**). Consistent with increased expression of pyroptotic pathway associated genes, snATAC-seq analysis showed increased open chromatin accessibility peaks of *CASP1*, *CASP9*, *IL1B* and *NLRP3*, in the β cell cluster of VMI organoids containing proinflammatory macrophages (**Figure 5I**). Upregulation of the pyroptotic pathway in the β cell cluster of VMI organoids containing proinflammatory macrophages was further confirmed by immunostaining using an antibody against CASP1 (**Figures 5J and 5K**). Apart from β cell pyroptosis, we did not observe β cell dedifferentiation in VMI organoids with pro-inflammatory macrophages (**Figure S5I**). Together, these data suggest that proinflammatory macrophages can induce β cell pyroptosis.

### Mechanistic studies identify pathways contributing to proinflammatory macrophage-mediated β cell pyroptosis

To determine the potential mechanisms by which proinflammatory macrophages induce β cell pyropotosis, we performed cell-cell interaction (cell-chat) analysis and focused on the interactions from macrophages to β cells. First, when comparing differential signaling from macrophages to β cells in VMI organoids containing proinflammatory macrophages to VMI organoids containing unstimulated macrophages, we identified four enhanced macrophage-to-β cell interaction pathways, including *TNFSF12*-*TNFRSF12A*, *SPP1*- *ITGAV*+*ITGB1*, *F11R*-*F11R* and *DSC2*-*DSG2* (**Figure 6A**). Next, we examined cell-cell interactions from macrophages to β cells in human islets exposed to CVB4 virus and also found increased communication probability of the *TNFSF12*-*TNFRSF12A* pathway (**Figure 6B**). Furthermore, the expression level of *TNFSF12* was increased in macrophages of human islets exposed to SARS-CoV-2 (**Figure 6C**). Immunostaining confirmed the increased expression of TNFSF12 in both human islets exposed to viruses and VMI organoids containing proinflammatory macrophages (**Figures S6A and S6B**). These data indicate that the TNFSF12-TNFRSF12A pathway might contribute to proinflammatory macrophage mediated β cell pyroptosis.

**Figure 6.**
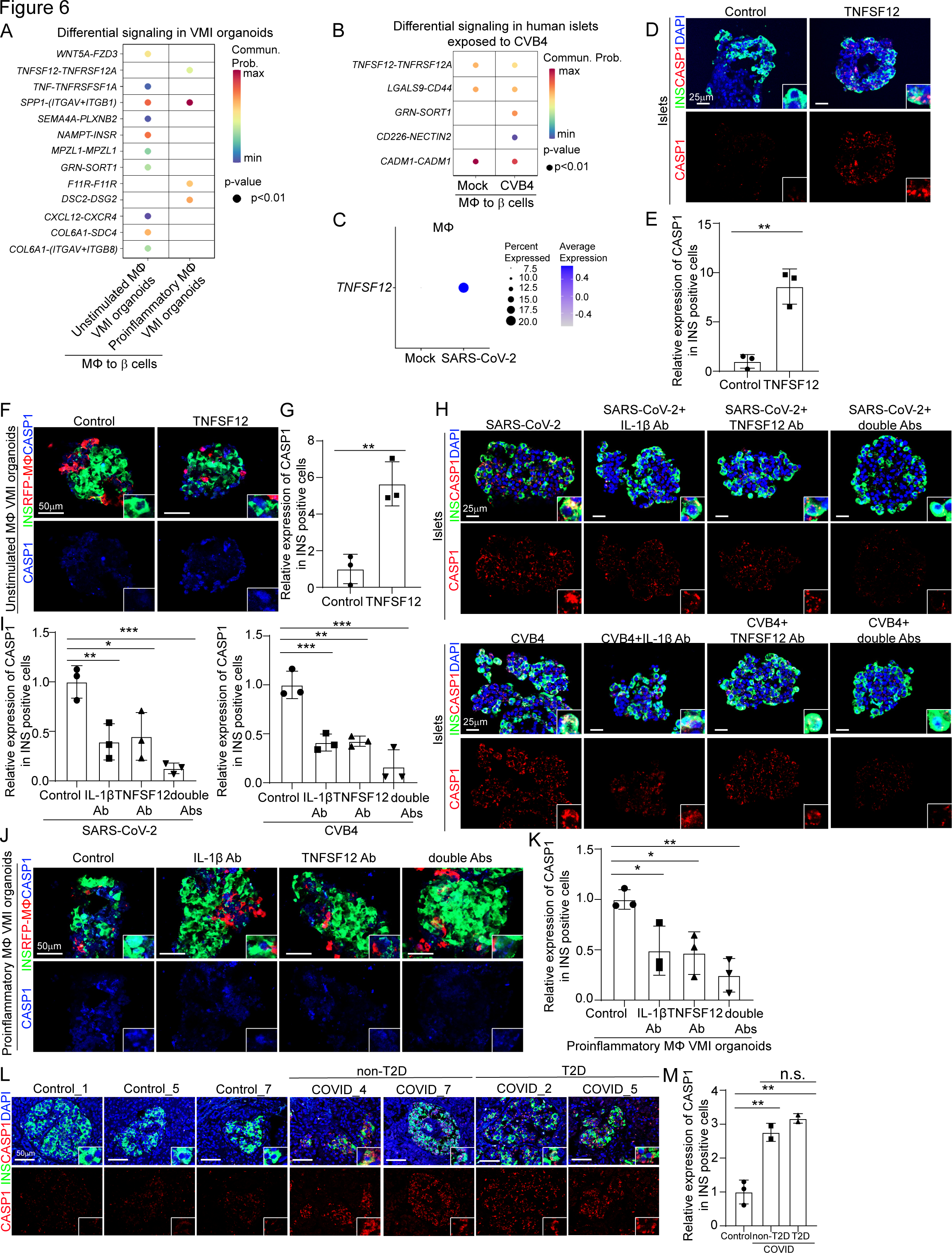
TNFSF12-TNFRSF12A as a candidate pathway that contributes to proinflammatory macrophage-mediated β cell pyroptosis. **(A)** Dot plot showed the differential signaling from macrophages to β cells in VMI organoids containing unstimulated or proinflammatory macrophages at day 7 after reaggregation. **(B)** Dot plot showed the differential signaling from macrophages to β cells in human islets exposed to mock or CVB4 virus (2x10^6^ PFU/ml). **(C)** Dot plot of the expression level of *TNFSF12* in macrophage of human islets exposed to mock or SARS-CoV-2 virus (MOI=1). **(D and E)** Confocal images (D) and quantification (E) of CASP1 expression in INS^+^ cells in control or 10 ng/ml TNFSF12 treated human islets. Scale bar= 25 µm. **(F and G)** Confocal images (F) and quantification (G) of CASP1 expression in INS^+^ cells in control or 10 ng/ml TNFSF12 treated VMI organoids at day 7 after reaggregation. Scale bar= 50 µm. **(H and I)** Confocal images (H) and quantification (I) of CASP1 expression in INS^+^ cells of SARS-CoV-2 (MOI=0.5) or CVB4 (2x10^6^ PFU/ml) exposed human islets treated with control, 10 µg/ml TNFSF12 blocking antibody, 5 µg/ml IL-1β blocking antibody or 10 µg/ml TNFSF12 + 5 µg/ml IL-1β blocking antibodies. Scale bar= 25 µm. **(J and K)** Confocal images (J) and quantification (K) of CASP1 expression in INS^+^ cells of VMI organoids containing proinflammatory macrophages at day 7 after reaggregation and treated with control, 10 µg/ml TNFSF12 blocking antibody, 5 µg/ml IL-1β blocking antibody or 10 µg/ml TNFSF12 + 5 µg/ml IL-1β blocking antibodies. Scale bar= 50 µm. **(L and M)** Confocal images (L) and quantification (M) of the CASP1 expression in INS^+^ cells of pancreas autopsy samples from control (N=3) and COVID-19 (N=4) subjects. The insert shows a high magnification of cells. Scale bar= 50 µm. N=3 independent biological replicates. Data was presented as mean ± STDEV. *P* values were calculated by unpaired two-tailed Student’s t test or one-way ANOVA with a common control. n.s., no significance; \**P* < 0.05, \*\**P* < 0.01, \*\*\**P* < 0.001. See also Figure S6.

To validate the role of TNFSF12-TNFRSF12A in proinflammatory macrophage mediated β cells pyroptosis, we treated human islets and VMI organoids containing unstimulated macrophages with TNFSF12 protein and detected increased CAPS1 expression in INS^+^ β cells in both cases (**Figures 6D-6G**). Next, we tested TNFSF12 neutralization antibody and found that it partially blocked β cell pyroptosis caused by SARS-CoV-2 or CVB4 exposure (**Figures 6H and 6I**), suggesting that other factors, might also contribute to this process. One candidate is IL-1β, which was detected in the supernatant of VMI organoids containing proinflammatory macrophages and macrophages of human islets exposed to SARS-CoV-2 or CVB4 **(Figures 2C, 2J, 5C and Figures S2J-2K, S3C-S3D)** and was reported to contribute to cell pyroptosis^67,68^. Indeed, IL-1β neutralization antibody partially blocked the increased β cell pyroptosis in human islets exposed to SARS-CoV-2 or CVB4. Furthermore, the combination of IL-1β and TNFSF12 neutralization antibodies showed add-on/synergistic effect to further decrease the CASP1 expression levels in human islets exposed to SARS-CoV-2 or CVB4 (**Figures 6H and 6I**) and VMI organoids with proinflammatory macrophages or VMI organoids exposed to SARS-CoV-2 or CVB4 (**Figures 6J-6K and Figures S6C-S6F**). Finally, we stained the pancreatic autopsy samples and confirmed the increased CASP1 expression in COVID-19 samples compared to control samples (**Figures 6L and 6M**). Moreover, the increased CASP1 expression was independent of T2D conditions (**Figures 6L and 6M**). GeoMx transcriptomic data also showed increased expression of pyroptosis-associated genes in ROIs of islets in COVID-19 samples compared to control samples (**Figure S6G**).

## Discussion

While several spatial transcriptomic analyses have been applied to study COVID-19 autopsy samples, they have focused on lung^69–71^, liver^72^, heart^73^, and placenta^74^. In this study, we used the GeoMx spatial transcriptomics and proteomics platform to comprehensively analyze changes in the immune cell composition and endocrine cell damage of COVID-19 pancreatic samples. Our findings revealed accumulation of proinflammatory macrophages in islets of COVID-19 samples, which highlights the critical role of macrophages in pathological changes observed in host tissues in COVID-19 patients. Previous study has shown that SARS-CoV2 induces a pro-fibrotic signature in monocytes, which includes CD163, a marker not expressed in homeostatic monocytes^75^. We also see an increase of CD163+ macrophages in islets of COVID-19 samples, which could be an indication of fibrogenic monocyte infiltration. Fibrosis might also play a role for pancreatitis, new onset diabetes and thus β cell damage^76^. Fibrotic alterations might be another potential driver of tissue dysfunction besides β cell pyroptosis. Surprisingly, in our spatial transcriptomics data, we didn’t see an increase of T cells in islets of COVID- 19 samples while both CD4 and CD8 T cells contribute to T1D development^77,78^. Upon thorough examination of published studies, no reports were identified regarding T cell infiltrations in the islets of COVID-19 samples. This underscores the need to impartially assess immune cell accumulation and expand the scope of investigation by examining additional COVID-19 pancreas samples.

In our study, we found that β cells in VMI organoids showed improved maturity. Islet vascular endothelial cells were reported to promote insulin production and secretion, as well as β-cell proliferation, survival, and maturation, by secreting a variety of growth factors, components of the extracellular matrix (ECM), and other molecules^79–81^. Macrophages exist in the pancreas from the embryonic stage onward. While the role of macrophages in islet morphogenesis is not well understood, various observations underscore their significance in the formation of the endocrine pancreas, especially in the development of β cells^82,83^.

While immune-mediated host damage is recognized as a critical factor in various diseases, there is a scarcity of suitable human in vitro models. Here, we constructed a hPSC-derived VMI organoid model, which allowed us to dissect molecular mechanisms of macrophage-mediated host damage. Through cell-cell interaction analysis, we found that proinflammatory macrophages induce β cell pyroptosis through the TNFSF12- TNFRSF12A pathway. Previous studies in the context of cholestasis demonstrated that bile acids induce TNFRSF12A expression, subsequently initiating hepatocyte pyroptosis through the NFκB/Caspase-1/GSDMD signaling pathway^84^. TNFSF12-TNFRSF12A pathway has been reported to contribute to the hepatocyte pyroptosis through NFκB/Caspase-1/GSDMD signaling^84^. Persistent TNFSF12-TNFRSF12A signaling has been implicated in the pathogenesis of numerous diseases, including atherosclerosis, ischemic stroke, rheumatoid arthritis (RA), and inflammatory bowel diseases^85,86^. Some of the TNFSF12-TNFRSF12A targeted therapeutic agents under development for these conditions^87^. Enavatuzumab, BIIB036 and RG7212, the humanized monoclonal antibodies targeting the TNFSF12-TNFRSF12A signaling, were tested in patients with tumors^88–90^. BIIB023 was also tested in patients with Rheumatoid Arthritis (NCT00771329) and lupus nephritis (NCT01499355) in clinical trials^91^. Here, we identified a previously unknown role of TNFSF12-TNFRSF12A in macrophages induced β cell pyroptosis.

Besides, we also explored the cell-cell interactions from β cells to immune cells (**Figure S6H**). β cell death constitutes a pathophysiological cornerstone in the natural progression of diabetes. Previous investigations into β cell death have primarily centered on apoptosis, necrosis, and autophagy. In this study, we uncovered a previously unknown mechanism in which proinflammatory macrophages induce β cell pyroptosis. An expanding body of research has linked β cell pyroptosis to diabetes^92, 93^. These findings suggest that macrophage-mediated β cell pyroptosis may contribute to the increased incidence of diabetes among COVID-19 patients.

## Limitations of study

In this study, we analyzed the pancreatic autopsy samples from non-COVID and COVID19 subjects. COVID-19 subjects might have experienced bed resting and starvation (ICU), which could have influenced the β cell phenotype, including insulin content. Additionally, the inherent variability/heterogeneity of studying pancreatic autopsy samples could pose analytical challenges in distinguishing genuine disease pathology or differences between human donors from experimental noise^94^. The modest sample size and potential confounders in the clinical samples could also be limitations in this study. In VMI organoids, we found that some endothelial cells can form small vessels. However, they cannot form intact blood vessels, which are likely required for monocyte infiltration into tissue. The vascular structure of VMI organoids is not fully functional yet, suggesting a need for further modification of the culture conditions. This might also be why we didn’t find obvious infiltration of monocytes into organoids containing endocrine cells and endothelial cells. Using this VMI organoid model, we observed that pro-inflammatory macrophage activation induced β cell death. However, we cannot distinguish whether the observed effects were derived from monocyte derived or tissue resident macrophages.

## ACKNOWLEDGMENTS

This work was supported by the National Institute of Diabetes, Digestive and Kidney Diseases (NIDDK, R01DK137517, and R01 DK124463, 1R01DK130454, S.C.), Department of Surgery, Weill Cornell Medicine (T.E., S.C.), and R01DK121072 Department of Medicine, Weill Cornell Medicine (R.E.S.), by S.C. and R.E.S. were supported as Irma Hirschl Trust Research Award Scholars. Human islets were received from the University of Pennsylvania human islet center with funding provided by the NIDDK supported Human Pancreas Analysis Program (HPAP) (https://hpap.pmacs.upenn.edu/citation) grants UC4 DK112217 to A.N.. Integrated Islet Isolation and Distribution Program (IIDP) NIH grants UC4DK098085 to A.N.. V.G. is a Weill Cornell Department of Medicine *Fund for the Future* awardee, supported by the Kellen Foundation. The authors would like to thank Didier Hober for kindly providing CVB4 virus. The authors would like to thank the Electron Microscopy & Histology services of the Weill Cornell Medicine Microscopy & Image Analysis Core and funds from an NIH Shared Instrumentation Grant (S10RR027699) for Shared Resources. The authors also thank Dr. Mike Drdos from National Human Genome Research Institute for his help on islet collection.

## AUTHOR CONTRIBUTIONS

S. C., R.E.S., L.Y. Q.J., S. C.J. P., and T.E. conceived and designed the experiments.

L. Y., Y.H., T.Z., X. D. T. L., J.J. V. N. d. S., C. C. R., and A. R. performed cell differentiation and immunostaining.

C. P., Y.S., and J. X. assisted with the library preparation.

J.G. performed SARS-CoV-2 infections.

C. L., W.W., and A.L. prepared the human islets and human pancreas samples.

R.E.S. provided autopsy sample of COVID-19 patients.

T. Z. and J. Z. performed the bioinformatics analyses.

## STAR*METHODS

Detailed methods are provided in the online version of this paper and include the following:

KEY RESOURCES TABLE

## RESOURCE AVAILABILITY

### Lead contact

Further information and requests for resources, reagents or codes should be directed to and will be fulfilled by the Lead Contact, Shuibing Chen.

### Materials availability

This study did not generate new unique reagents.

### Data and code availability

scRNA-seq, snATAC-seq and RNA-seq data have been deposited at GEO and are publicly available as of the date of publication. Accession numbers are listed in the key resources table. All original code has been deposited at Github and is publicly available as of the date of publication. DOI is listed in the key resources table. Any additional information required to reanalyze the data reported in this paper is available from the lead contact upon request.

## METHOD DETAILS

### Human studies

Pancreas tissues from COVID-19 samples were provided by the Weill Cornell Medicine Department of Pathology using protocols approved by the Tissue Procurement Facility of Weill Cornell Medicine. Experiments using samples from human subjects were conducted in accordance with local regulations and with the approval of the IRB at the Weill Cornell Medicine. The autopsy samples were collected under protocol 20-04021814. For GeoMx RNA and protein analysis, seven COVID19 human pancreas samples were deceased upon tissue acquisition and were provided from Weill Cornell Medicine as fixed samples. The control human pancreas samples were obtained from the Human Islet Core at the University of Pennsylvania. The pancreatic organs were obtained from the organ procurement organization under the United Network for Organ Sharing. The organs were kept in the University of Wisconsin solution at 4°C before the tissue samples biopsies. The freshly dissected tissues (<3mm thick) were fixed with 10% formalin for 8 hours at room temperature. The tissue samples were rinsed with running tap water for 5 min then through 80% and 95% alcohol for 1 hour each, followed with 2 rinses of 100% alcohol for 1 hour each for dehydration. The tissues were cleared in xylene 3 times for 1 hour each. The tissues were immersed in paraffin 3 times for 1 hour each before being embedded in a paraffin block. The paraffin-embedded tissue blocks were sectioned at 5 μm thickness on a microtome and floated in a 40°C water bath containing distilled water. The sections were transferred onto glass slides which were suitable for immunohistochemistry and the slides were dried at room temperature before use.

### hPSC maintenance and pancreatic differentiation

*INS^GFP/W^* MEL-1 cells were used to generate pancreatic endocrine cells using a previously reported strategy^95^. In brief, *INS^GFP/W^* MEL-1 cells were cultured on Matrigel-coated 6-well plates in StemFlex medium (Gibco Thermo Fisher) and maintained at 37℃ with 5% CO2. At stage 1-day 1, cells were exposed to basal RPMI 1640 medium supplemented with 1× Glutamax (Thermo Fisher Scientific), 50 μg/mL Normocin, 100 ng/mL Activin A (R&D systems), and 3 μM of CHIR99021 (GSK3β inhibitor 3, Cayman Chemical) for 24 hours. At stage 1-day 2 and 3, the medium was changed to basal RPMI 1640 medium supplemented with 1× Glutamax, 50 μg/mL Normocin, 0.2% fetal bovine serum (FBS, Corning), 100 ng/mL Activin A for 2 days. At stage 2-day 4 and 5, the resulting definitive endoderm cells were cultured in MCDB131 medium (Thermo Fisher Scientific) supplemented with 1.5 g/L sodium bicarbonate, 1× Glutamax, 10 mM glucose (Sigma Aldrich) at final concentration, 2% bovine serum albumin (BSA, Lampire), 0.25 mM L-ascorbic acid (Sigma Aldrich) and 50 ng/ml of fibroblast growth factor 7 (FGF-7, Peprotech) to acquire primitive gut tube. At stage 3-day 6 and day 7, cells were induced to differentiate to posterior foregut in MCDB 131 medium supplemented with 2.5 g/L sodium bicarbonate, 1× Glutamax, 10 mM glucose at final concentration, 2% BSA, 0.25 mM L-ascorbic acid, 50 ng/ml of FGF-7, 1 μM Retinoic acid (RA; Sigma Aldrich), 100 nM LDN193189 (LDN, Axon Medchem), 1:200 ITS-X (Thermo Fisher Scientific), 200 nM TPPB (Tocris Bioscience) and 0.25 μM SANT-1 (Sigma Aldrich) for 2 days. At stage 4-day 8-day 10, cells were differentiated to pancreatic endoderm in MCDB 131 medium supplemented with 2.5 g/L sodium bicarbonate, 1× Glutamax, 10 mM glucose at final concentration, 2% BSA, 0.25 mM L-ascorbic acid, 2 ng/ml of FGF-7, 0.1 μM RA, 200 nM LDN193189, 1:200 ITS-X, 100 nM TPPB and 0.25 μM SANT-1 for 3 days. At stage 5-day 11-day 13, cells were differentiated to pancreatic endocrine precursors in MCDB 131 medium supplemented with 1.5 g/L sodium bicarbonate, 1× Glutamax, 20 mM glucose at final concentration, 2% BSA, 0.05 μM RA, 100 nM LDN, 1:200 ITS-X, 0.25 μM SANT-1, 1 mM T3 hormone (Sigma Aldrich), 10 μM ALK5 inhibitor II (Cayman Chemical), 10 μM zinc sulfate heptahydrate (Sigma Aldrich) and 10 μg/ml of heparin (Sigma Aldrich) for 3 days. At day 14, cells were exposed to MCDB 131 medium supplemented with 1.5 g/L sodium bicarbonate, 1× Glutamax, 20 mM glucose at final concentration, 2% BSA, 100 nM LDN193189, 1:200 ITS-X, 1 μM T3, 10 μM ALK5 inhibitor II, 10 μM zinc sulfate, 10 μg/ml of heparin, 100 nM gamma secretase inhibitor XX (Millipore) for 7 days. Then, cells were exposed to MCDB 131 medium supplemented with 1.5 g/L sodium bicarbonate, 1× Glutamax, 20 mM glucose at final concentration, 2% BSA, 1:200 ITS-X, 1 μM T3, 10 μM ALK5 inhibitor II, 10 μM zinc sulfate heptahydrate, 10 μg/ml of heparin, 1 mM N-acetyl cysteine (Sigma Aldrich), 10 μM Trolox (Millipore), 2 μM R428 (MedchemExpress) for another 7-15 days. The medium was subsequently refreshed every day.

### hPSC differentiation toward endothelial cells

To derive endothelial cells from hPSCs, we optimized a previously reported strategy^96^. Briefly, H1 hESCs were passaged onto Matrigel-coated 6-well plates in StemFlex medium. Before differentiation, we infected H1 hESCs with lentivirus carrying ETV2. After two days selection with 1 µg/ml puromycin and 1 day recovery in StemFlex medium, hESCs will be switched to StemDiff APEL medium (STEMCELL Technologies) with 6 µM CHIR99021 for 2 days. Then, cells were cultured in StemDiff APEL medium with an additional of 25 ng/ml BMP-4, 10 ng/ml bFGF and 50 ng/ml VEGF (R&D Systems) for another two days. On day 4, cells were dissociated with Accutase (Innovative Cell Technologies) and reseeded onto p100 culture dishes in EC Growth Medium MV2 (Promocell) with an additional 50 ng/ml VEGF for 4-6 days. Finally, endothelial cells were generated and passaged every 3-5 days in EC Growth Medium MV2 with an additional 50 ng/ml VEGF. Before coculture as organoids or non-coculture as control, hPSCs-derived endothelial cells were purified by magnetic sorting using anti- CD31 (PECAM1) beads.

### hPSCs differentiation towards macrophages

H9 hESCs expressing RFP (RFP-H9) were differentiated using a previously reported protocol^97^. RFP-H9 cells were dissociated with ReLeSR (STEMCELL Technologies) as small clusters onto Matrigel-coated 6-well plates at low density. The day after passaging, cells were cultured in IF9S medium supplemented with 50 ng/ml BMP-4, 15 ng/ml Activin A and 1.5 µM CHIR99021. After 2 days, medium was refreshed with IF9S medium supplemented with 50 ng/ml VEGF, 50 ng/ml bFGF, 50 ng/ml SCF (R&D Systems) and 10 µM SB431542 (Cayman Chemical). On day 5 and 7, medium was changed into IF9S medium supplemented with 50 ng/ml IL- 6 (R&D Systems), 10 ng/ml IL-3 (R&D Systems), 50 ng/ml VEGF, 50 ng/ml bFGF, 50 ng/ml SCF and 50 ng/ml TPO (R&D Systems). On day 9, cells were dissociated with TrypLE (Life Technologies) and resuspended in IF9S medium supplemented with 50 ng/ml IL-6, 10 ng/ml IL-3 and 80 ng/ml M-CSF (R&D Systems). On day 13, medium was changed to IF9S medium supplemented with 50 ng/ml IL-6, 10 ng/ml IL-3 and 80 ng/ml M-CSF. On day 15, monocytes can be collected at this stage for further experiments. Otherwise, monocytes can be collected and plated on FBS-coated plates in IF9S medium supplemented with 80 ng/ml M-CSF to generate macrophages. IF9S medium was prepared according to previous publication^97^. All differentiation steps were cultured under normoxic conditions at 37 ℃, 5% CO2. Before coculture as organoids or non-coculture as control, hPSCs-derived macrophages were purified by magnetic sorting using anti-CD14 beads.

### GeoMx transcriptomic and protein assays

Human control and COVID-19 pancreas samples were prepared as FFPE slides and applied to the NanoString GeoMx® Digital Spatial Profiler platform according to the manufacturer’s instructions. In brief, slides from FFPE embedded pancreatic autopsy samples were prepared two weeks before experiments. Insulin (INS), Pan-ck (Pan Cytokeratin) and nuclear dye (TOTO™-3 Iodide) were used as morphology markers for selecting ROIs. We selected 6 ROIs in human islet areas, 3 ROIs in exocrine area and 3 ROIs in ductal area for each pancreas sample. The protein assays and transcriptomic assays were performed using adjacent sides. Data analysis was performed on GeoMx DSP software.

### Construction of VMI organoids

The VMI organoids were constructed with hPSC- derived pancreatic endocrine cells, endothelial cells, and macrophages. Briefly, endocrine cells were dissociated with Accutase (Innovative Cell Technologies) at Day 16- 19, macrophages were dissociated with Accutase after day 19 of the differentiation procedure, and endothelial cells were dissociated with Trypsin 0.25% EDTA (THERMO FISHER) after day 10 of the differentiation procedure. The dissociated single cells were reaggregated with approximately 70-80% pancreatic endocrine cells, 10-20% endothelial cells, and approximately 2-5% macrophages in VMI organoid culture medium containing 80% pancreatic endocrine cells’ stage 6 medium (MCDB 131 medium supplemented with 1.5 g/L sodium bicarbonate, 1× Glutamax, 20 mM glucose at final concentration, 2% BSA, 100 nM LDN193189, 1:200 ITS-X, 1 μM T3, 10 μM ALK5 inhibitor II, 10 μM zinc sulfate, 10 μg/ml of heparin, 100 nM gamma secretase inhibitor XX) plus 20% endothelial cells’ medium (EC Growth Medium MV2 with an additional 50 ng/ml VEGF) using low-attach U plates. 48 hours later, the cells self-assembled into organoids. Subsequently, the medium was changed every two days.

### Human islets

The pancreatic organs were obtained from the local organ procurement organization under the United Network for Organ Sharing (UNOS). The islets were isolated in the Human Islet Core at the University of Pennsylvania following the guidelines of Clinical Islet Transplantation consortium protocol^98^. Briefly, the pancreas was digested following intraductal injection of Collagenase & Neutral Protease in Hanks’ balanced salt solution. Liberated islets were then purified on continuous density gradients (Cellgro/Mediatech) using the COBE 2991 centrifuge and cultured in CIT culture media and kept in a humidified 5% CO2 incubator.

### Cell Lines

HEK293T (human [*Homo sapiens*] fetal kidney) and Vero E6 (African green monkey [*Chlorocebus aethiops*] kidney) were obtained from ATCC). Cells were cultured in Dulbecco’s Modified Eagle Medium (DMEM) supplemented with 10% FBS and 100 I.U./mL penicillin and 100 μg/mL streptomycin. All cell lines were incubated at 37°C with 5% CO2.

### SARS-CoV-2 Viruses and infection

SARS-CoV-2, isolate USA-WA1/2020 was obtained from World Reference Center for Emerging Viruses and Arboviruses located at University of Texas Medical Branch via the CDC. Vero E6 cells (ATCC) served as the culture system for SARS-CoV-2 propagation, utilizing EMEM with a supplement of 10% FCS, 1 mM Sodium Pyruvate, and 10 mM HEPES (citation). All work involving live SARS-CoV-2 was performed in the CDC/USDA-approved BSL-3 facility at Aaron Diamond AIDS Research Center located at Columbia University. The Aaron Diamond AIDS Research Center’s BSL-3 facility at Columbia University prepared the SARS-CoV-2 WA1 strain, subsequently stored at -70°C. Infection assays on human islets or hESCs-derived VMI organoids were conducted in culture medium at specified multiplicity of infections (MOIs) and incubated at 37°C. Post-infection, at predetermined hours post-infection (hpi), the cells underwent triple PBS washes and a 60-minute fixation in 4% formaldehyde at room temperature. Culture medium alone served as control.

### CVB4 Viruses and infection

The aliquots of CVB4 E2, the diabetogenic strain of coxsackievirus B4 virus were provided by Didier Hober and were then stored frozen at −80°C. Human islets or hESCs-derived VMI organoids were infected with CVB4 E2 at 2 × 10^6^ PFU/ml (2 × 10^4^ PFU/organoid)^99^. Human islets or VMI organoids were then incubated in a humidified incubator at 37°C with 5% CO2 at predetermined hours post-infection (hpi), the cells underwent triple PBS washes and a 60-minute fixation in 4% formaldehyde at room temperature. Culture medium alone served as control.

### Immunohistochemistry

Tissues were fixed overnight in 4% buffered formalin and transferred to 30% sucrose before being snap-frozen in O.C.T (Fisher Scientific, Pittsburgh, PA). Live organoids in culture were directly fixed in 4% paraformaldehyde for 30 min, followed with 60 min of permeabilization and blocking in PBS supplemented with 0.2% Triton X-100 and 5% horse serum. For immunofluorescence, cells or tissue sections were stained with primary antibodies at 4°C overnight and secondary antibodies at RT for 1h. The information for primary antibodies and secondary antibodies were provided in Table S3. Nuclei were counterstained by DAPI.

### Single-cell RNA-seq data analysis of human islets upon SARS-CoV-2 or CVB4 infection

The 10X libraries were sequenced on the Illumina NovaSeq6000 sequencer with paired end reads (28 bp for read 1 and 90 bp for read 2). Subsequently, the sequencing data were primarily analyzed using the 10X cellranger pipeline v6.1.1 in a two-step process. In the initial step, cellranger *mkfastq* demultiplexed the samples and generated fastq files. In the subsequent step, cellranger *count* aligned the fastq files to a customized reference genome, extracting a gene expression UMI counts matrix for each library. The customized reference genome was constructed by integrating the 10X pre-built human reference GRCh38-2020-A, the SARS-CoV-2 virus genome, and the CVB4 virus genome using the cellranger *mkref*. The two virus genomes were obtained from the NCBI Nucleotide database with accession numbers NC_045512.2 (SARS-CoV-2) and AF311939.1 (CVB4).

We applied several filtering criteria, excluding cells with fewer than 500 or more than 6000 detected genes, cells with fewer than 1000 or more than 60000 detected transcripts, and cells with mitochondrial gene content exceeding 15%. Subsequently, we employed a deconvolution strategy^100^ for normalizing gene expression UMI counts, utilizing the R scran package (v.1.14.1). Specifically, we initiated the process by pre-clustering cells with the *quickCluster* function. We then computed size factors per cell within each cluster, rescaled these factors by normalization between clusters using the *computeSumFactors* function, and normalized the UMI counts per cell by the size factors, followed by a logarithmic transform using the *normalize* function. We further normalized UMI counts across samples using the *multiBatchNorm* function in the R batchelor package (v1.2.1). We employed solo^101^ v0.6 to identify doublets, which were subsequently excluded from the downstream analysis.

We identified highly variable genes using the *FindVariableFeatures* function in the R Seurat package v3.1.0, selecting the top 3000 variable genes after excluding mitochondria genes, ribosomal genes, viral genes, and dissociation-related genes. The list of dissociation-related genes, originally built on mouse data, was converted to human ortholog genes using Ensembl BioMart. Cells from multiple samples were aligned based on their mutual nearest neighbors (MNNs)^102^ using the *fastMNN* function in the R batchelor package v1.2.1. This involved performing principal component analysis (PCA) on the highly variable genes and then correcting the principal components (PCs) according to their MNNs. We chose the corrected top 50 PCs for downstream visualization and clustering analysis.

Uniform Manifold Approximation and Projection (UMAP) dimensional reduction were executed using the *RunUMAP* function in the R Seurat package, with the number of neighboring points set to 30 and the training epochs set to 4000. Cells were clustered into thirteen clusters by constructing a shared nearest neighbor graph and grouping cells of similar transcriptome profiles using the *FindNeighbors* function and *FindClusters* function (resolution set to 0.2) in the R Seurat package. After reviewing the clusters, we merged them into nine clusters representing acinar cells, α cells, β cells, δ cells, ductal cells, mesenchymal cells, PP cells, immune cells, and endothelial cells for further analysis. Marker genes for the merged nine clusters were identified by performing differential expression analysis between cells inside and outside each cluster using the *FindMarkers* function in the R Seurat package. The expressions of cell type markers within each cell population were depicted through violin plots, utilizing the *VlnPlot* function in the R Seurat package. The expression of CVB4-polyprotein were visualized either through UMAP plot, employing the Seurat *DimPlot* function, or via jitter plot created with R ggplot2 package v3.2.1.

To assess cell death associated pathways within varied cell types following SARS-CoV- 2 infection, we compared gene expressions in α cells, β cells, δ cells, mesenchymal cells and endothelial cells between mock and SARS-CoV-2 infected conditions using the Wilcoxon rank-sum test via the *FindMarkers* function in the R Seurat package. Subsequently, we ordered the genes based on log2 fold change and performed gene set enrichment analysis on cell death associated pathways using the *GSEA* function in the R clusterProfiler^103^ package v4.6.2. The expressions of pyroptosis pathway associated genes in β cells under mock and SARS-CoV-2 infected conditions were visualized using the *DotPlot* function in the R Seurat package. The expressions of HLA genes and autoantigen associated genes in β cells under mock, SARS-CoV-2 infection or CVB4 infection conditions were represented using violin plots generated with the *VlnPlot* function in the R Seurat package.

To investigate the immune cell population, we extracted immune cells and performed a sub-clustering analysis. Highly variable genes were identified using the *FindVariableFeatures* function in the R Seurat package, and the top 3000 variable genes were selected, excluding mitochondria genes, ribosomal genes, viral genes and dissociation-related genes. Cells from multiple samples were aligned using the *fastMNN* function in the R batchelor package, as described above. The top 50 corrected PCs were selected for UMAP dimensional reduction using the *RunUMAP* function in the R Seurat package, with the number of neighboring points set to 30 and training epochs setting to 200. The immune cell population was clustered into seven clusters using the *FindNeighbors* function and *FindClusters* function (resolution set to 0.8) in the R Seurat package. After reviewing these clusters, we merged them into five clusters representing macrophages, dendritic cells, immune progenitor cells, T cells and B cells.

UMAP and violin plots were generated to illustrate the cell clusters and highlight expressions of selected genes using the R ggplot2 package v3.2.1. Dot plots were generated to show gene expression changes in the mock and infected conditions using the *DotPlot* function in the R Seurat package.

### Single-cell RNA-seq data analysis of VMI organoids

The 10X scRNA-seq libraries underwent sequencing on the Illumina NovaSeq6000 sequencer with pair-end reads (28 bp for read 1 and 90 bp for read 2). Subsequently, the sequencing data were primarily analyzed using the 10X cellranger pipeline v7.1.0 in a two-step process. In the initial step, cellranger *mkfastq* demultiplexed the samples and generated fastq files. In the subsequent step, cellranger *count* aligned the fastq files to the 10X pre-built human reference GRCh38-2020-A reference, extracting a gene expression UMI counts matrix for each library.

Several filtering criteria were applied, excluding cells with fewer than 300 or more than 9000 detected genes, cells with fewer than 600 or more than 75000 detected transcripts, and cells with mitochondrial gene content exceeding 10%. Doublet cells in each sample were identified, assuming a doublet rate 0.8% per 1000 recovered cells, as reported by 10X Genomics, using the R DoubletFinder^104^ package v2.0.3. The doublet cells were subsequently excluded from downstream analysis.

We employed a deconvolution strategy^100^ for normalizing gene expression UMI counts, utilizing the R scran (v.1.22.1), scuttle (v1.4.0) and batchelor (v1.10.0) packages. The process involved pre-clustering cells with the *quickCluster* function in the R scran package, computing size factors per cell within each cluster, rescaling these factors by normalization between clusters using the *computeSumFactors* function in the R scran package, normalizing the UMI counts per cell by the size factors, followed by a logarithmic transform using the *logNormCounts* function in the R scuttle package. Further normalization of UMI counts across samples was performed using the *multiBatchNorm* function in the R batchelor package. Cells from multiple samples were aligned based on their MNNs using the *quickCorrect* function in the R batchelor package. This involved identifying highly variable genes, performing PCA on the highly variable genes and then correcting the PCs according to their MNNs. The corrected top 50 PCs were chosen for downstream clustering analysis. The corrected gene expression values on variable genes were reconstructed based on the corrected PCs and were used for coembeding scRNA- seq and snATAC-seq data.

UMAP dimensional reduction were executed using the *RunUMAP* function in the R Seurat package^105^ av4.1.0, with the number of neighboring points set to 30 and the training epochs set to 500. Cells were clustered into fourteen clusters by constructing a shared nearest neighbor graph and grouping cells of similar transcriptome profiles using the *FindNeighbors* function and *FindClusters* function (resolution set to 0.5) in the R Seurat package. After reviewing the clusters, we merged them into nine clusters representing acinar cells, α cells, β cells, δ cells, ductal cells, endocrine progenitor cells, endothelial cells, macrophages and proliferation cells for further analysis.

DE analysis was performed on β cells between VMI organoids and non-coculture cells, and between VMI organoids with and without proinflammatory macrophages using the Wilcoxon rank-sum test via the *FindMarkers* function in the R Seurat package. Volcano plots were generated to illustrate DE genes using the R ggplot2 package v3.4.2. Dot plots were generated to show gene expression changes in different clusters or conditions using the *DotPlot* function in the R Seurat package. Pie charts were utilized to visualize cell type compositions within VMI organoids using the R ggplot package v3.4.2.

To determine the mechanisms by which proinflammatory macrophages induce β cell pyropotosis, we conducted cell-cell interaction analysis between macrophage and β cell populations using the R CellChat^106^ package v1.5.0. Bubble plots were generated to illustrate the communication probabilities mediated by ligand-receptor pairs in between macrophage and β cell populations using the *netVisual_bubble* function.

### Single-nuclear ATAC-seq data analysis of VMI organoids

The 10X snATAC-seq libraries underwent sequencing on the Illumina NovaSeq6000 sequencer with pair-end reads (51bp for read 1 and 51bp for read 2). Subsequently, the sequencing data were primarily analyzed using the 10X cellranger-atac pipeline v2.1.0 in a two-step process. In the initial step, cellranger-atac *mkfastq* demultiplexed the samples and generated fastq files. In the subsequent step, cellranger-atac *count* aligned the fastq files to the 10X pre-built GRCh38 2020-A-2.0.0 reference, performed peak calling, and extracted a barcoded and aligned fragment file for each library.

We ultilized the R Signac^107^ package v1.10.0 to analyze snATAC-seq data. Specifically, we created a common set of peaks across all samples using the *reduce* function and generated a peaks x cell matrix for each sample by quantifying the common peaks using the *FeatureMatrix* function. We applied several filtering criteria, excluding cells with fewer than 3000 or more than 30000 peaks detected, cells with fewer than 20% of reads in peaks, cells with more than 5% of reads in blacklist regions, cells with the ratio of mononucleosomal to nucleosome-free fragments greater than 4, and cells with TSS enrichment score smaller than 3. Term frequency-inverse document frequency (TF-IDF) normalization was performed using the *RunTFIDF* function. We selected the top-ranked peaks using the *FindTopFeatures* function and ran singular value decomposition (SVD) to obtain latent semantic indexing (LSI) components using the *RunSVD* function. The top 50 LSI components, excluding the first LSI component, were used for downstream clustering analysis.

We classified cells into the nine cell types based on clustering results from scRNA-seq data. This was achieved by quantifying gene expression activity from the snATAC-seq data using the *GeneActivity* function, identifying anchors between scRNA-seq and snATAC-seq data using the *FindTransferAnchors* function, and transferring the cell clustering labels from scRNA-seq to snATAC-seq data using the *TransferData* function.

We co-embeded the scRNA-seq and snATAC-seq cells in the same UMAP plot. This was done by imputing gene expressions for the snATAC-seq cells based on the corrected gene expression values from the scRNA-seq cells using the *TransferData* function, merging cells from scRNA-seq and snATAC-seq, scaling the expression values and performing PCA using the *ScaleData* and RunPCA functions, and selecting the top 30 PCs for UMAP dimensional reduction using the *RunUMAP* function with the number of neighboring points setting to 30 and training epochs setting to 500.

UMAP plots were generated to illustrate the cell clusters using the R ggplot2 package v3.4.2. Aggregated chromatin accessibility signals were visualized for multiple groups of cells within a given genomic region using the *CoveragePlot* function. Chromatin accessibility signal for individual cells were visualized using the *TilePlot* function. Pie charts were utilized to visualize cell type compositions within VMI organoids using the R ggplot package v3.4.2.

### Bulk RNA-seq data analysis

The libraries underwent sequencing with single-end 50 bps on the Illumina NovaSeq6000 sequencer. Raw sequencing reads in BCL format were processed through bcl2fastq 2.20 (Illumina) for FASTQ conversion and demultiplexing. After trimming the adaptors with cutadapt v1.18, the sequencing reads were mapped to the human GRCh37 reference by STAR^108^ av2.5.2b. Read counts per gene were extracted using HTSeq-count v0.11.2^109^, and normalized through a regularized log transformation with the DESeq2 package v1.26.0^110^.

### RNA-Seq

Total RNA was extracted in TRIzol (Invitrogen) and DNase I treated using Directzol RNA Miniprep kit (Zymo Research) according to the manufacturer’s instructions. RNAseq libraries of polyadenylated RNA were prepared using the TruSeq RNA Library Prep Kit v2 (Illumina) or TruSeq Stranded mRNA Library Prep Kit (Illumina) according to the manufacturer’s instructions. cDNA libraries were sequenced using an Illumina NextSeq 500 platform. The resulting single end reads were checked for quality (FastQC v0.11.5) and processed using the Digital Expression Explorer 2 (DEE2)^111^ workflow. Adapter trimming was performed with Skewer (v0.2.2)^112^. Further quality control done with Minion, part of the Kraken package^113^. The resultant filtered reads were mapped to human reference genome GRCh38 using STAR aligner^108^ and gene-wise expression counts generated using the “-quantMode GeneCounts” parameter. BigWig files were generated using the bamCoverage function in deepTools2 (v.3.3.0)^114^. After further filtering and quality control, R package edgeR^115^ was used to calculate RPKM and Log2 counts per million (CPM) matrices as well as perform differential expression analysis. Heatmap was generated using online tool: http://www.heatmapper.ca/.

## QUANTIFICATION AND STATISTICAL ANALYSIS

N=3 independent biological replicates were used for all experiments unless otherwise indicated. *P*-values were calculated by unpaired two-tailed Student’s t-test or one way ANOVA with a common control unless otherwise indicated. n.s. indicates a non-significant difference. \**p*<0.05, \*\**p*<0.01 and \*\*\**p*<0.001.

**Supplemental Video 1.** VMI organoids at day 14 after reaggregation. β cells: INS-GFP; macrophages: RFP; and endothelial cells: Far Red. **Related to Figure 3**.

**Supplemental Video 2.** VMI organoids at day 14 after reaggregation stained with antibodies against INS (Green), CD68 (Red) and PECAM1 (CD31, Blue). **Related to Figure 3**.

**Supplemental Video 3.** VMI organoids at day 14 after reaggregation stained with antibodies against INS (Green), CD68 (Red), GCG (Blue) and PECAM1 (CD31, Gray). **Related to Figure 3**.

**Supplemental Video 4.** VMI organoids at day 14 after reaggregation stained with antibodies against INS (Green), CD68 (Red), SST (Blue) and PECAM1 (CD31, Gray). **Related to Figure 3**.

**Supplemental Video 5.** VMI organoids at day 14 after reaggregation stained with antibodies against INS (Green), CD68 (Red), NKX6.1 (Blue) and PECAM1 (CD31, Gray). **Related to Figure 3**.

**Supplemental Video 6.** Live imaging of calcium signaling in VMI organoids upon high glucose stimulation. High glucose: 20 mM D-glucose. Each frame was captured every 500 ms. **Related to Figure 3**.

**Supplemental Video 7.** Live imaging of VMI organoids at day 7 after reaggregation exposed with with CVB4 virus (2x10^6^PFU/ml). β cells: INS-GFP, macrophages: RFP. **Related to Figure 3**.

**Supplemental Video 8.** VMI organoids at day 7 after reaggregation containing unstimulated macrophages stained with antibodies against INS (Green), CD68 (Red), PECAM1 (CD31, Gray) and DAPI (Blue). **Related to Figure 5**.

**Supplemental Video 9.** VMI organoids at day 7 after reaggregation containing proinflammatory macrophages stained with antibodies against INS (Green), CD68 (Red), PECAM1 (CD31, Gray) and DAPI (Blue). **Related to Figure 5**.

