## Supplemental Figures for "Human Vascularized Macrophage-Islet Organoids to Model Immune-Mediated Pancreatic β cell Pyroptosis upon Viral Infection"

### Figure S1

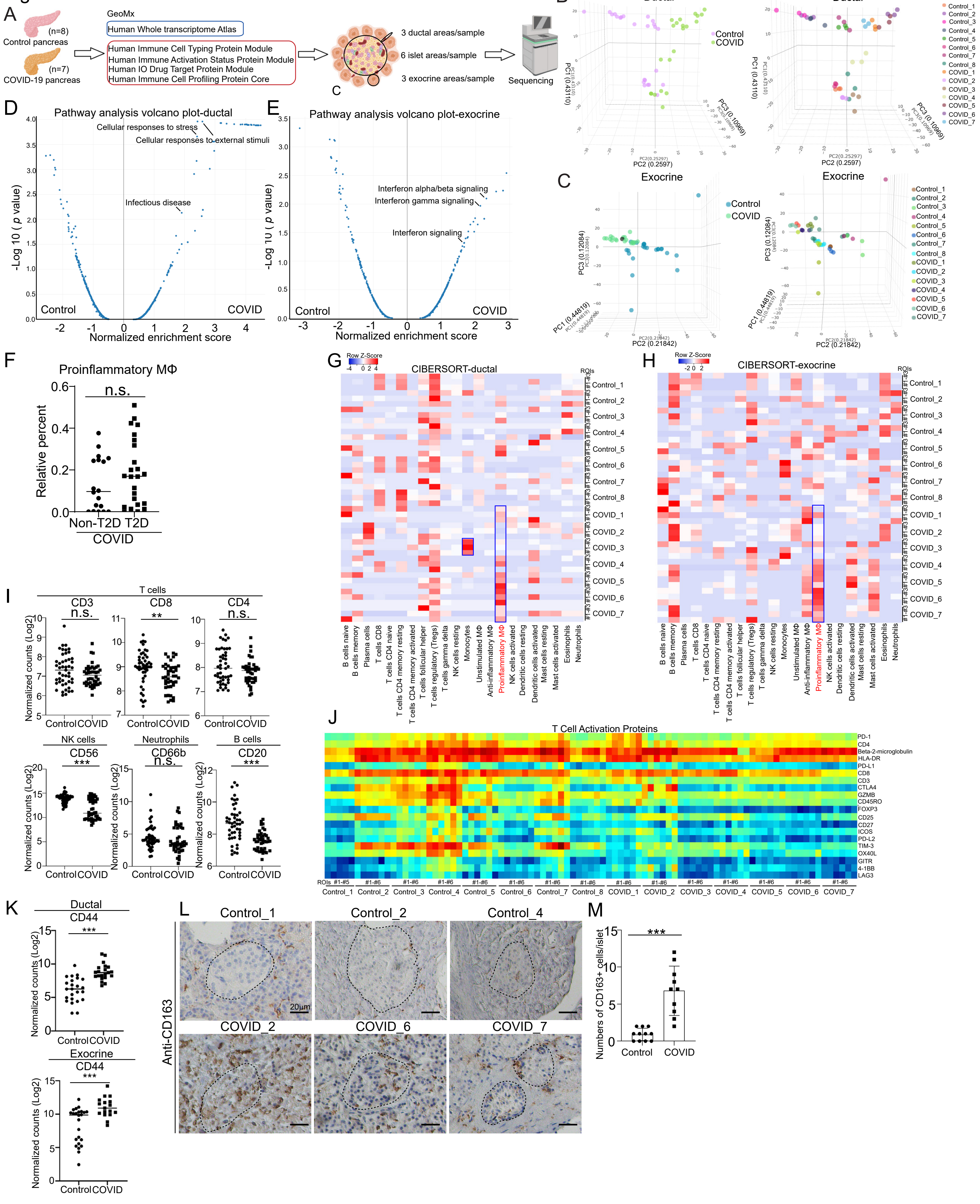

A
GeoMx
(n=8)
Control pancreas
(n=7)
COVID-19 pancreas
Human Whole transcriptome Atlas
Human Immune Cell Typing Protein Module
Human Immune Activation Status Protein Module
Human IO Drug Target Protein Module
Human Immune Cell Profiling Protein Core
3 ductal areas/sample
6 islet areas/sample
3 exocrine areas/sample
Sequencing
B
Ductal
Control
COVID
C
Exocrine
D
Pathway analysis volcano plot-ductal
Cellular responses to stress
Cellular responses to external stimuli
Infectious disease
Normalized enrichment score
E
Pathway analysis volcano plot-exocrine
Interferon alpha/beta signaling
Interferon gamma signaling
Interferon signaling
F
Proinflammatory MΦ
n.s.
Relative percent
Non-T2D T2D
COVID
G
CIBERSORT-ductal
Row Z-Score
H
CIBERSORT-exocrine
I
T cells
CD3
CD8
CD4
NK cells
CD56
Neutrophils
CD66b
B cells
CD20
Normalized counts (Log2)
Control COVID
J
T Cell Activation Proteins
PD-1
CD4
Beta-2-microglobulin
HLA-DR
PD-L1
CD8
CD3
CTLA4
GZMB
CD45RO
FOXP3
CD25
CD27
ICOS
PD-L2
TIM-3
OX40L
GITR
4-1BB
LAG3
K
Ductal
CD44
Exocrine
CD44
L
Control_1
Control_2
Control_4
COVID_2
COVID_6
COVID_7
Anti-CD163
20μm
M
Numbers of CD163+ cells/islet
Control
COVID

#### SUPPLEMENTAL INFORMATION

**Figure S1. GeoMx spatial transcriptomics and protein analysis of pancreatic autopsy samples from control and COVID-19 subjects. Related to Figure 1.**

**(A)** Schematic representation of GeoMx spatial transcriptomics and protein analysis.

**(B and C)** 3D PCA plots of GeoMx whole transcriptome sequencing data from ROIs of human ductal (B) and exocrine (C) areas of COVID-19 (N=7) and control (N=8) pancreatic autopsy samples.

**(D and E)** Volcano plot highlighting the pathways enriched in transcriptome sequencing data from ROIs of human ductal (D) and exocrine (E) areas of COVID-19 (N=7) and control (N=8) pancreatic autopsy samples.

**(F)** Relative percent of proinflammatory macrophages from the CIBERSORT analysis of immune cells (LM22) using the GeoMx whole transcriptome sequencing data of human islet areas in non-T2D or T2D COVID-19 pancreatic autopsy samples. Each dot represents one count in each ROI.

**(G)** Heatmap of the CIBERSORT analysis of immune cells (LM22) using the GeoMx whole transcriptome sequencing data from ROIs of human ductal areas of COVID-19 (N=7) and control (N=8) pancreatic autopsy samples.

**(H)** Heatmap of the CIBERSORT analysis of immune cells (LM22) using the GeoMx whole transcriptome sequencing data from ROIs of human exocrine areas of COVID-19 (N=7) and control (N=8) pancreatic autopsy samples.

**(I)** Normalized counts (Log2) of immune cell markers, including CD3, CD8 and CD4 for T cells, CD56 for NK cells, CD66b for neutrophils and CD20 for B cells, in ROIs of the islet areas of control (N=8) and COVID-19 (N=7) pancreatic autopsy samples. Each dot represents one count in each ROI.

**(J)** Heatmap of the proteins related to T cell activation examined by GeoMx protein assay from ROIs of human islet areas of COVID-19 (N=7) and control (N=8) pancreatic autopsy samples.

**(K)** Normalized counts (Log2) of CD44 in the ductal or exocrine areas of control (N=8) and COVID-19 (N=7) pancreatic autopsy samples. Each dot represents one count in each ROI.

**(L and M)** Immunohistochemistry staining (L) and quantification (M) of CD163 in COVID-19 (N=3) and control (N=3) pancreatic autopsy samples. Dotted lines encircled the regions of the islets. Scale bar=20 μm.

*P* values were calculated by unpaired two-tailed Student's t test. n.s., no significance, $^{**}P < 0.01$, $^{***}P < 0.001$.

Figure S2

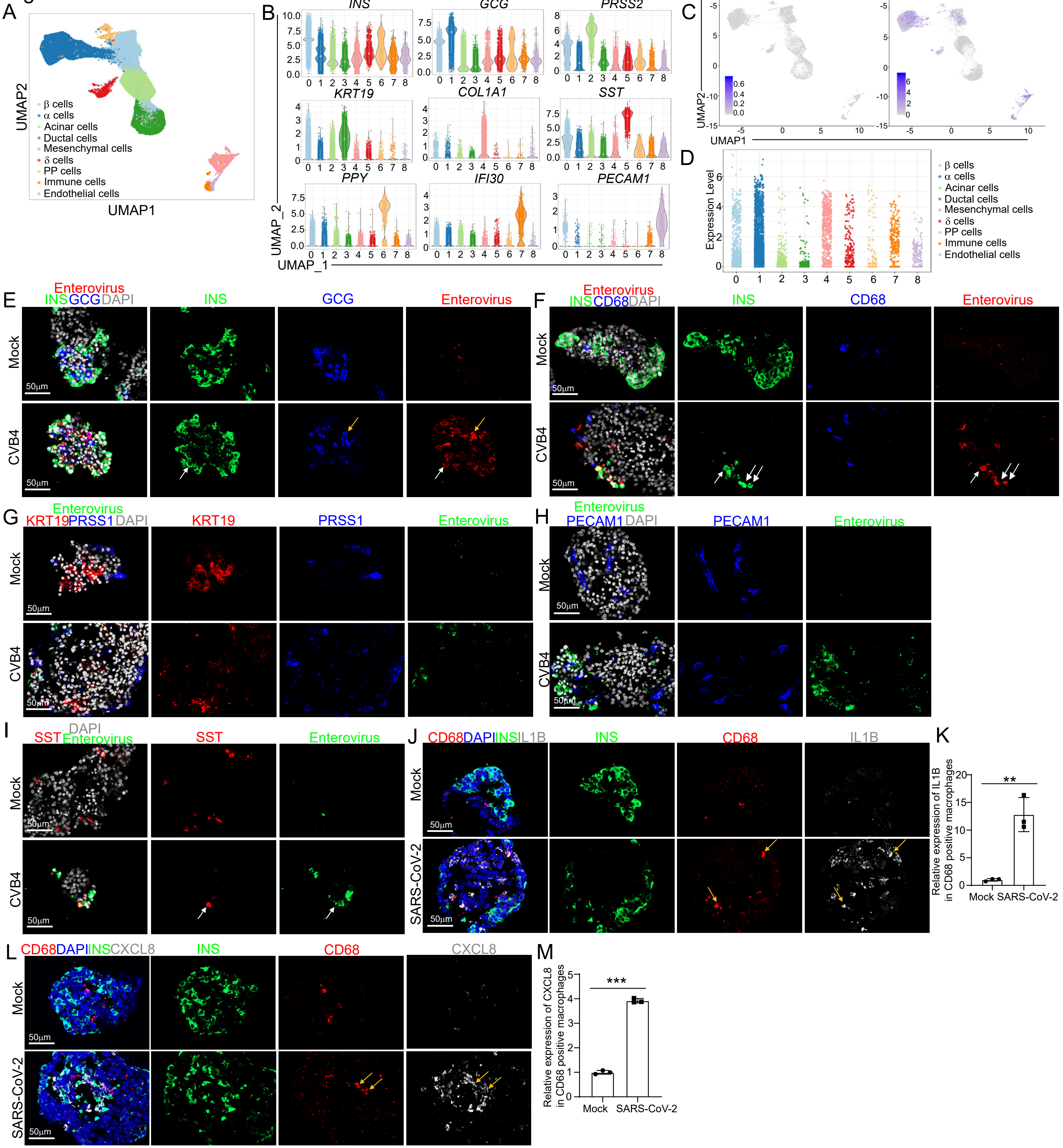
A
UMAP2
UMAP1
β cells
α cells
Acinar cells
Ductal cells
Mesenchymal cells
δ cells
PP cells
Immune cells
Endothelial cells
B
INS
GCG
PRSS2
KRT19
COL1A1
SST
PPY
IFI30
PECAM1
UMAP_2
UMAP_1
C
Mock-polyprotein
CVB4--polyprotein
UMAP2
UMAP1
D
Expression Level
E
Enterovirus
INS GCG DAPI
INS
GCG
Enterovirus
Mock
CVB4
50μm
F
Enterovirus
INS CD68 DAPI
INS
CD68
Enterovirus
Mock
CVB4
G
Enterovirus
KRT19 PRSS1 DAPI
KRT19
PRSS1
Enterovirus
H
Enterovirus
PECAM1 DAPI
PECAM1
Enterovirus
I
DAPI
SST Enterovirus
SST
Enterovirus
J
CD68 DAPI INS IL1B
INS
CD68
IL1B
SARS-CoV-2
K
Relative expression of IL1B in CD68 positive macrophages
**
Mock
SARS-CoV-2
L
CD68 DAPI INS CXCL8
INS
CD68
CXCL8
M
Relative expression of CXCL8 in CD68 positive macrophages
***
Mock
SARS-CoV-2

**Figure S2. CVB4 infects human islets and induces activation of proinflammatory macrophages. Related to Figure 2.**

**(A)** UMAP of human islets exposed to mock, SARS-CoV-2 (MOI=1) or CVB4 ($2x10^6$ PFU/ml) viruses.

**(B)** Violin plot of cell markers of each cell population, including *INS, GCG, PRSS2, KRT19*, *COL1A1*, *SST*, *PPY*, *IFI30*, and *PECAM1*.

**(C)** UMAP showed the expression of CVB4-*polyprotein* in human islets exposed to mock or CVB4 ($2x10^6$ PFU/ml) virus.

**(D)** Jitter plot showed the expression of CVB4-*polyprotein* in human islets exposed to mock or CVB4 ($2x10^6$ PFU/ml) virus.

**(E)** Confocal images of Enterovirus (CVB4) antigen expression in $INS^+$ β cells, $GCG^+$ α cells of human islets exposed to mock or CVB4 ($2x10^6$ PFU/ml). The white arrows highlight the co-localization of INS and Enterovirus antigen. The yellow arrows highlight the co-localization of GCG and Enterovirus antigen. Scale bar= 50 μm.

**(F)** Confocal images of Enterovirus (CVB4) antigen expression in $INS^+$ β cells, $CD68^+$ macrophages of human islets exposed to mock or CVB4 ($2x10^6$ PFU/ml). The white arrows highlight the co-localization of INS and Enterovirus antigen. Scale bar= 50 μm.

**(G)** Confocal images of Enterovirus (CVB4) antigen expression in $KRT19^+$ ductal cells, $PRSS1^+$ acinar cells of human islets exposed to mock or CVB4 ($2x10^6$ PFU/ml). Scale bar= 50 μm.

**(H)** Confocal images of Enterovirus (CVB4) antigen expression in $PECAM1^+$ endothelial cells of human islets exposed to mock or CVB4 ($2x10^6$ PFU/ml). Scale bar= 50 μm.

**(I)** Confocal images of Enterovirus (CVB4) antigen expression in $SST^+$ δ cells of human islets exposed to mock or CVB4 ($2x10^6$ PFU/ml). The white arrows highlight the co-localization of SST and Enterovirus antigen. Scale bar= 50 μm.

**(J and K)** Confocal images (J) and quantification (K) of IL1B expression in $CD68^+$ macrophages of human islets exposed to mock or SARS-CoV-2 (MOI=0.5). The yellow arrows highlight the co-localization of CD68 and IL1B. Scale bar= 50 μm.

**(L and M)** Confocal images (L) and quantification (M) of CXCL8 expression in $CD68^+$ macrophages of human islets exposed to mock or SARS-CoV-2 (MOI=0.5). The yellow arrows highlight the co-localization of CD68 and CXCL8. Scale bar= 50 μm.

*P* values were calculated by unpaired two-tailed Student's t test. $^{**}P < 0.01$, $^{***}P < 0.001$.

Figure S3

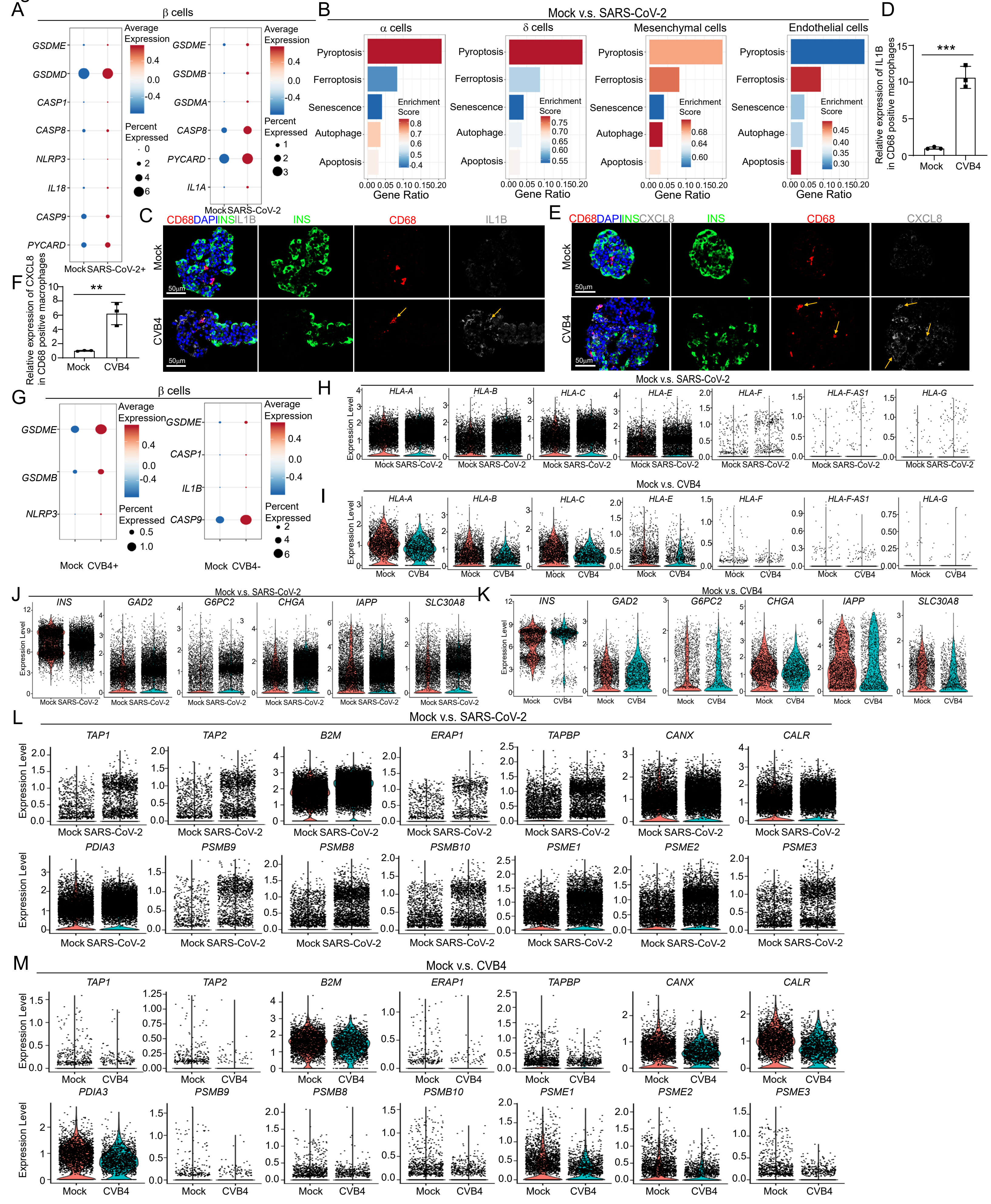

A
β cells
GSDME
GSDMD
CASP1
CASP8
NLRP3
IL18
CASP9
PYCARD
Mock SARS-CoV-2+
Average Expression
Percent Expressed
GSDME
GSDMB
GSDMA
CASP8
PYCARD
IL1A
Mock SARS-CoV-2
B
Mock v.s. SARS-CoV-2
α cells
δ cells
Mesenchymal cells
Endothelial cells
Pyroptosis
Ferroptosis
Senescence
Autophage
Apoptosis
Enrichment Score
Gene Ratio
C
CD68DAPIINSIL1B
INS
CD68
IL1B
Mock
CVB4
50μm
D
Relative expression of IL1B in CD68 positive macrophages
***
Mock
CVB4
E
CD68DAPIINSCXCL8
INS
CD68
CXCL8
F
Relative expression of CXCL8 in CD68 positive macrophages
**
G
β cells
GSDME
GSDMB
NLRP3
Mock CVB4+
GSDME
CASP1
IL1B
CASP9
Mock CVB4-
H
Mock v.s. SARS-CoV-2
HLA-A
HLA-B
HLA-C
HLA-E
HLA-F
HLA-F-AS1
HLA-G
Expression Level
I
Mock v.s. CVB4
J
Mock v.s. SARS-CoV-2
INS
GAD2
G6PC2
CHGA
IAPP
SLC30A8
K
Mock v.s. CVB4
L
Mock v.s. SARS-CoV-2
TAP1
TAP2
B2M
ERAP1
TAPBP
CANX
CALR
PDIA3
PSMB9
PSMB8
PSMB10
PSME1
PSME2
PSME3
M
Mock v.s. CVB4

**Figure S3. Single cell RNA-seq analysis of human islets upon CVB4 or SARS-CoV-2 exposure. Related to Figure 2.**

**(A)** Dot plot analysis of pyroptosis pathway associated genes in virus$^+$ β cell cluster and virus$^-$ β cell cluster of human islets upon mock or SARS-CoV-2 exposure (MOI=1).

**(B)** Pathway enrichment analysis of cell death pathways in α, δ, mesenchymal and endothelial cell clusters of human islets exposed to mock or SARS-CoV-2 (MOI=1).

**(C and D)** Confocal images (C) and quantification (D) of IL1B expression in CD68$^+$ macrophages of human islets exposed to mock or CVB4 ($2x10^6$ PFU/ml). The yellow arrows highlight the co-localization of CD68 and IL1B. Scale bar= 50 μm.

**(E and F)** Confocal images (E) and quantification (F) of CXCL8 expression in CD68$^+$ macrophages of human islets exposed to mock or CVB4 ($2x10^6$ PFU/ml). The yellow arrows highlight the co-localization of CD68 and CXCL8. Scale bar= 50 μm.

**(G)** Dot plot analysis of pyroptosis pathway associated genes in virus$^+$ β cell cluster and virus$^-$ β cell cluster of human islets upon mock or CVB4 ($2x10^6$ PFU/ml).

**(H)** Violin plot of the expression of *HLA* genes in the β cell cluster of human islets exposed to mock or SARS-CoV-2 (MOI=1).

**(I)** Violin plot of the expression of *HLA* genes in the β cell cluster of human islets exposed to mock or CVB4 (2x$10^6$ PFU/ml).

**(J)** Violin plot of the expression of autoantigen genes in the β cell cluster of human islets exposed to mock or SARS-CoV-2 (MOI=1).

**(K)** Violin plot of the expression of autoantigen genes in the β cell cluster of human islets exposed to mock or CVB4 (2x$10^6$ PFU/ml).

**(L)** Violin plot of the expression of antigen presentation associated genes in the β cell cluster of human islets exposed to mock or SARS-CoV-2 (MOI=1).

**(M)** Violin plot of the expression of antigen presentation genes in the β cell cluster of human islets exposed to mock or CVB4 (2x$10^6$ PFU/ml).

*P* values were calculated by unpaired two-tailed Student's t test. $^{**}P < 0.01$, $^{***}P < 0.001$.

Figure S4

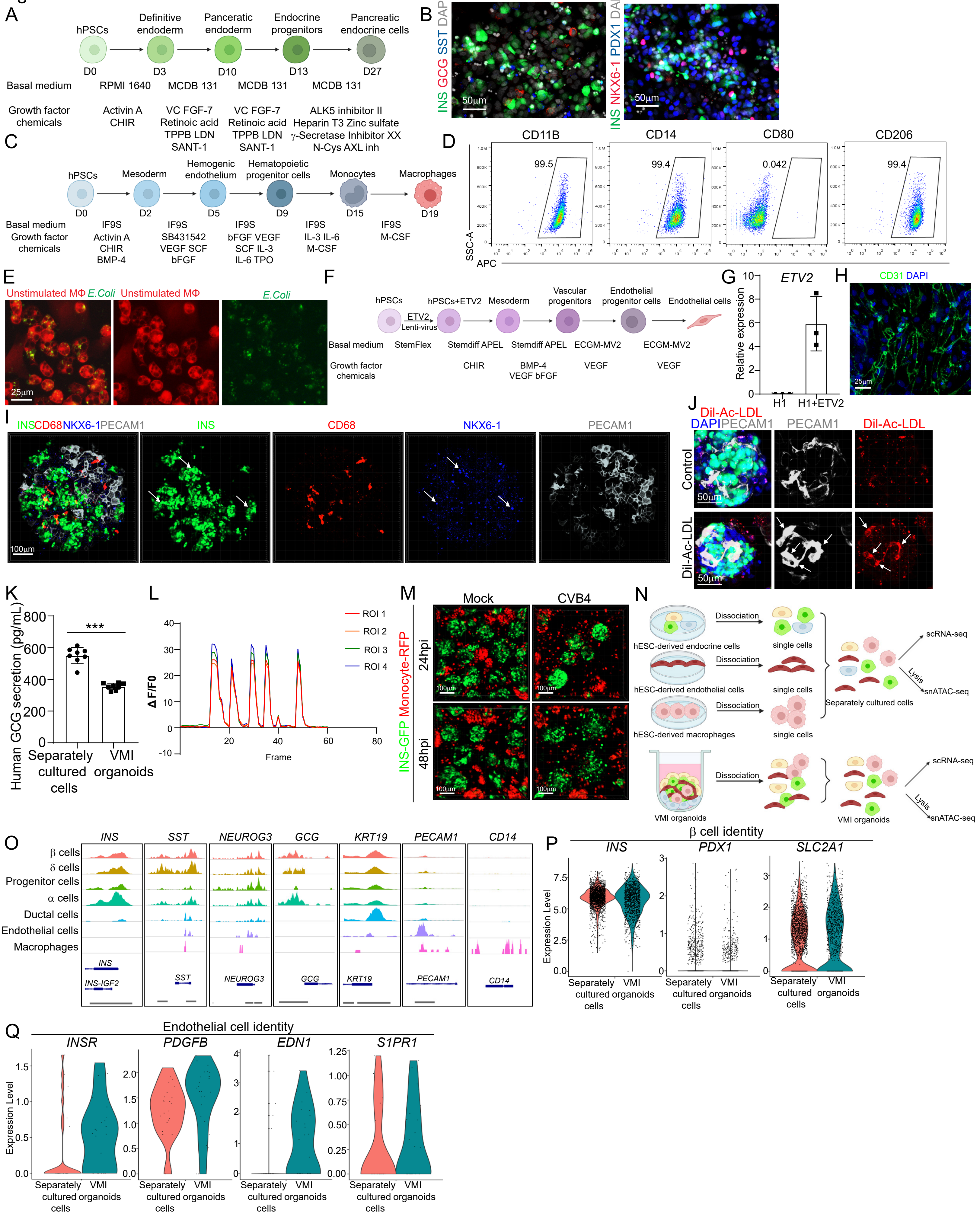

A
hPSCs
Definitive endoderm
Pancreatic endoderm
Endocrine progenitors
Pancreatic endocrine cells
D0
D3
D10
D13
D27
Basal medium
RPMI 1640
MCDB 131
MCDB 131
MCDB 131
Growth factor chemicals
Activin A CHIR
VC FGF-7 Retinoic acid TPPB LDN SANT-1
VC FGF-7 Retinoic acid TPPB LDN SANT-1
ALK5 inhibitor II Heparin T3 Zinc sulfate γ-Secretase Inhibitor XX N-Cys AXL inh
B
INS GCG SST DAPI
INS NKX6-1 PDX1 DAPI
50μm
C
hPSCs
Mesoderm
Hemogenic endothelium
Hematopoietic progenitor cells
Monocytes
Macrophages
D0
D2
D5
D9
D15
D19
Basal medium
Growth factor chemicals
IF9S Activin A CHIR BMP-4
IF9S SB431542 VEGF SCF bFGF
IF9S bFGF VEGF SCF IL-3 IL-6 TPO
IF9S IL-3 IL-6 M-CSF
IF9S M-CSF
D
CD11B
CD14
CD80
CD206
99.5
99.4
0.042
99.4
SSC-A
APC
E
Unstimulated MΦ E.Coli
Unstimulated MΦ
E.Coli
25μm
F
hPSCs
hPSCs+ETV2
Mesoderm
Vascular progenitors
Endothelial progenitor cells
Endothelial cells
ETV2 Lenti-virus
Basal medium
StemFlex
Stemdiff APEL
Stemdiff APEL
ECGM-MV2
ECGM-MV2
Growth factor chemicals
CHIR
BMP-4 VEGF bFGF
VEGF
VEGF
G
ETV2
Relative expression
H1
H1+ETV2
H
CD31 DAPI
25μm
I
INS CD68 NKX6-1 PECAM1
INS
CD68
NKX6-1
PECAM1
100μm
J
Dil-Ac-LDL DAPI PECAM1
PECAM1
Dil-Ac-LDL
Control
Dil-Ac-LDL
50μm
K
Human GCG secretion (pg/mL)
***
Separately cultured cells
VMI organoids
L
ROI 1
ROI 2
ROI 3
ROI 4
ΔF/F0
Frame
M
Mock
CVB4
24hpi
48hpi
INS-GFP Monocyte-RFP
100μm
N
hESC-derived endocrine cells
hESC-derived endothelial cells
hESC-derived macrophages
Dissociation
single cells
Separately cultured cells
scRNA-seq
Lysis
snATAC-seq
VMI organoids
O
INS
SST
NEUROG3
GCG
KRT19
PECAM1
CD14
β cells
δ cells
Progenitor cells
α cells
Ductal cells
Endothelial cells
Macrophages
INS-IGF2
P
β cell identity
INS
PDX1
SLC2A1
Expression Level
Separately cultured cells
VMI organoids
Q
Endothelial cell identity
INSR
PDGFB
EDN1
S1PR1
Expression Level
Separately cultured cells
VMI organoids

**Figure S4. Construction and characterization of hPSC-derived VMI organoids. Related to Figure 3 and Figure 4.**

**(A)** Schematic illustration of directed differentiation of hPSCs to pancreatic endocrine cells. At day 16, we detected INS-GFP$^{+}$ cells. Then, early stage 6 cells (day 16-day19) were co-cultured with macrophages and endothelial cells or culture separately for 7-14 days.

**(B)** Confocal images of hPSC-derived pancreatic endocrine cells stained with antibodies against INS, GCG, NKX6-1, PDX1 and SST. Scale bar= 50 μm.

**(C)** Schematic illustration of directed differentiation of hESCs to macrophages.

**(D)** Flow cytometry analysis of hPSC-derived unstimulated macrophages stained with antibodies against CD11B, CD14, CD80 and CD206.

**(E)** Confocal images of hPSC-derived macrophages engulfing GFP labeled *E.Coli*. Macrophages: RFP; *E.Coli: Green*. Scale bar= 25 μm.

**(F)** Schematic illustration of directed differentiation of hPSCs to endothelial cells.

**(G)** qRT-PCR analysis to examine the expression level of *ETV2* in H1 hPSCs following forced expression of ETV2 or control. Data was normalized to β-actin.

**(H)** Confocal images of hPSC-derived endothelial cells stained with antibodies against PECAM1 (CD31) and DAPI. Scale bar= 25 μm.

**(I)** Composite Z-stack confocal images of VMI organoids at day 14 after reaggregation stained with antibodies against INS, CD68, NKX6-1 and PECAM1 (CD31). The white arrows highlight the co-localization of INS and NKX6-1. Scale bar= 100 μm.

**(J)** Confocal images of VMI organoids stained with antibodies against PECAM1 (CD31) and Dil-Ac-LDL. Scale bar= 50 μm.

**(K)** ELISA assay showed the secretion of GCG in VMI organoids and separately cultured endocrine cells.

**(L)** Quantification of calcium signaling in VMI organoids upon high glucose stimulation. High glucose: 20 mM D-glucose. Each frame was captured every 500ms.

**(M)** Live cell imaging of VI organoids and monocytes exposed to mock or CVB4 ($2x10^6$ PFU/ml) at 24 hpi and 48 hpi. Scale bar= 100 μm.

**(N)** Schematic illustration of the sample preparation for scRNA-seq and snATAC-seq.

**(O)** Chromatin accessibility signals of cell markers for each cluster as analyzed using snATAC-seq.

(**P)** Violin plot analysis of β cell associated genes in β cell cluster of VMI organoids at day 7 after reaggregation and separately cultured cells as analyzed by scRNA-seq.

(**Q)** Violin plot analysis of endothelial cell associated genes in endothelial cell cluster of VMI organoids at day 7 after reaggregation and separately cultured cells as analyzed by scRNA-seq.

N=3 independent biological replicates. Data was presented as mean ± STDEV. ***$P$ < 0.001.

Figure S5

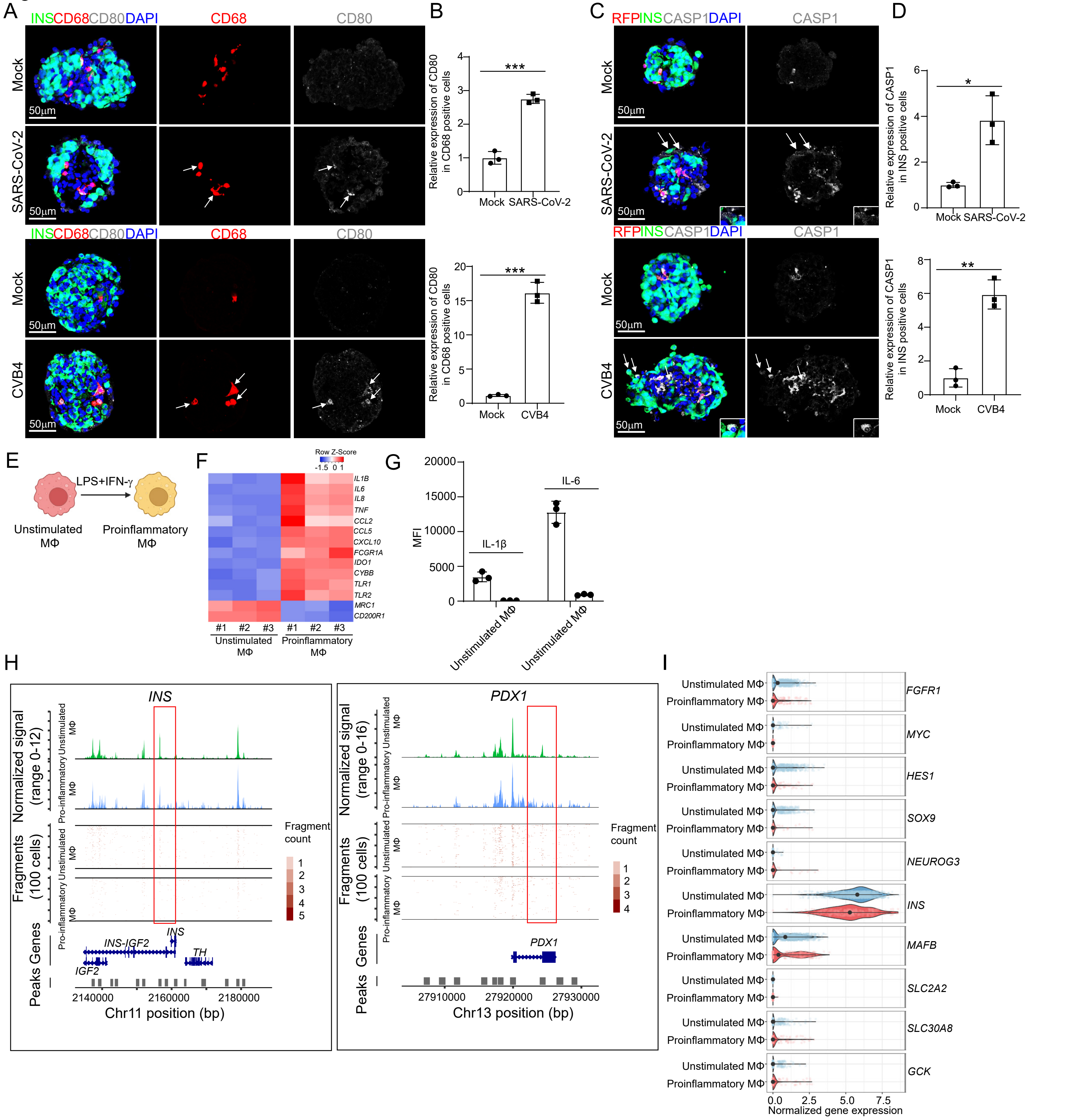
A
INSCD68CD80DAPI
CD68
CD80
Mock
SARS-CoV-2
CVB4
50μm
B
Relative expression of CD80 in CD68 positive cells
***
Mock
SARS-CoV-2
CVB4
C
RFPINSCASP1DAPI
CASP1
D
Relative expression of CASP1 in INS positive cells
*
**
E
LPS+IFN-γ
Unstimulated MΦ
Proinflammatory MΦ
F
Row Z-Score
IL1B
IL6
IL8
TNF
CCL2
CCL5
CXCL10
FCGR1A
IDO1
CYBB
TLR1
TLR2
MRC1
CD200R1
#1 #2 #3
Unstimulated MΦ
Proinflammatory MΦ
G
MFI
IL-1β
IL-6
Unstimulated MΦ
H
INS
PDX1
Normalized signal (range 0-12)
Normalized signal (range 0-16)
Fragments (100 cells)
Fragment count
Peaks Genes
INS-IGF2
IGF2
TH
Chr11 position (bp)
Chr13 position (bp)
I
FGFR1
MYC
HES1
SOX9
NEUROG3
INS
MAFB
SLC2A2
SLC30A8
GCK
Normalized gene expression

**Figure S5. Activation of proinflammatory macrophages and β cell pyroptosis were detected in hPSC-derived VMI organoids exposed to viruses. Related to Figure 5.**

**(A and B)** Confocal images (A) and quantification (B) of hPSC-derived VMI organoids exposed to viruses or mock conditions stained with antibodies against INS, CD68 and CD80 (SARS-CoV-2: MOI=0.5; CVB4: $2x10^6$ PFU/ml). Scale bar= 50 μm. The white arrows highlight the $CD68^+CD80^+$ cells.

**(C and D)** Confocal images (C) and quantification (D) of hPSC-derived VMI organoids exposed to viruses or mock conditions stained with antibodies against INS and CASP1 (SARS-CoV-2: MOI=0.5; CVB4: $2x10^6$ PFU/ml). Scale bar= 50 μm. The white arrows highlight the $INS^+CASP1^+$ cells.

**(E)** Schematic illustration of the stimulation of macrophages to proinflammatory macrophages.

**(F)** Heatmap showing the expression of macrophage associated genes in hPSC-derived macrophages with or without 2 days treatment with 100 ng/ml LPS and 20 ng/ml IFN-γ.

(**G)** The secretion of cytokines, including IL-1β and IL-6 in the supernatant of hPSC-derived macrophages with or without 2 days treatment with 100 ng/ml LPS and 20 ng/ml IFN-γ.

**(H)** Chromatin accessibility signals of the β cluster of VMI organoids at day 7 after reaggregation containing unstimulated macrophages or proinflammatory macrophages as analyzed using snATAC-seq. The normalized signal shows the averaged frequency of sequenced DNA fragments within a genomic region. The fragment shows the frequency of sequenced fragments within a genomic region for individual cells.

(**I)** Jitter plot analysis of β cell dedifferentiation associated genes in β cell cluster of VMI organoids with unstimulated macrophages or proinflammatory macrophages at day 7 after reaggregation as analyzed by scRNA-seq.

N=3 independent biological replicates. Data was presented as mean ± STDEV. $^{*}P < 0.05$, $^{**}P < 0.01$, $^{***}P < 0.001$.

Figure S6

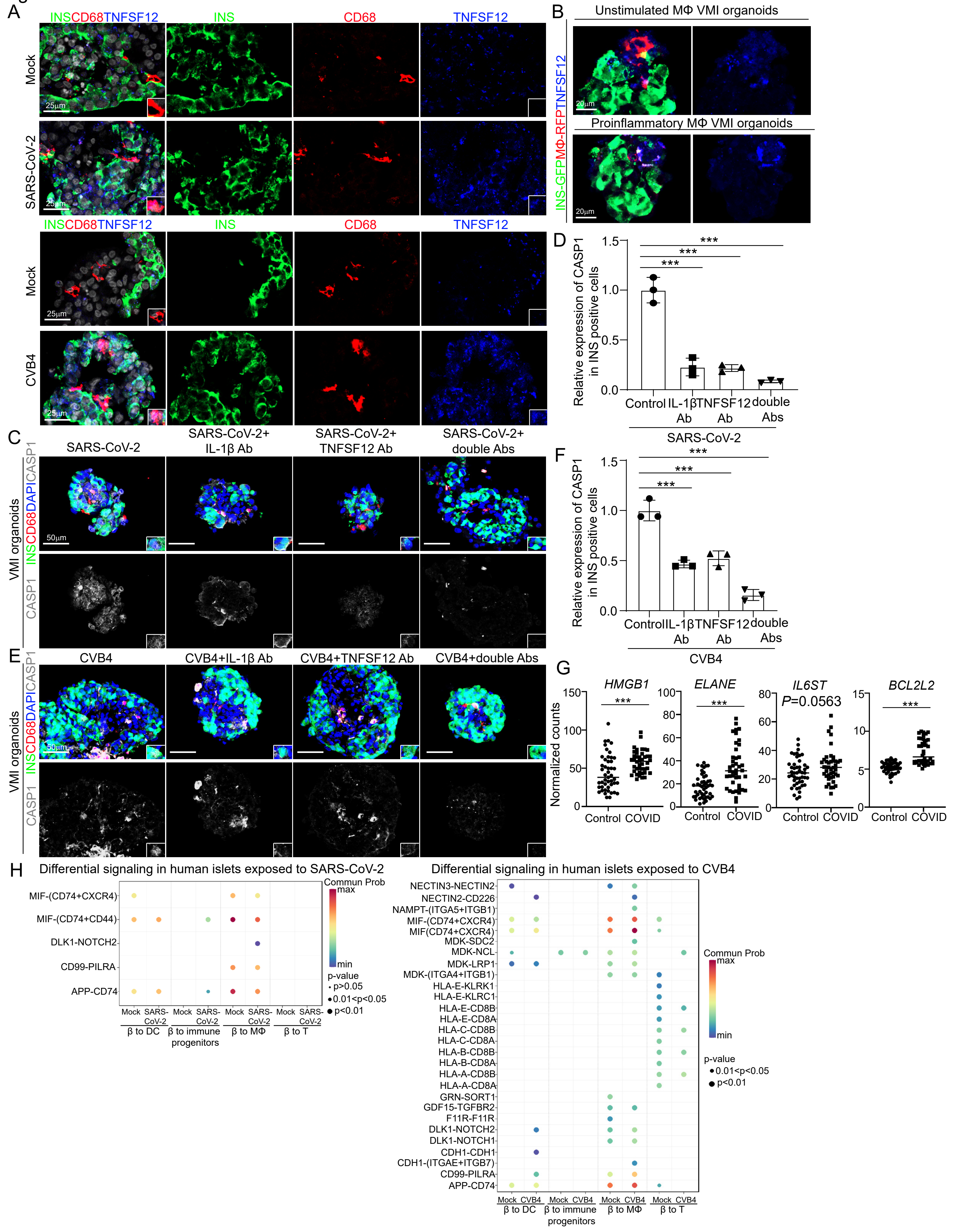
A
INSCD68TNFSF12
INS
CD68
TNFSF12
Mock
SARS-CoV-2
CVB4
25μm
B
Unstimulated MΦ VMI organoids
Proinflammatory MΦ VMI organoids
INS-GFPMΦ-RFPTNFSF12
20μm
C
SARS-CoV-2
SARS-CoV-2+ IL-1β Ab
SARS-CoV-2+ TNFSF12 Ab
SARS-CoV-2+ double Abs
VMI organoids
INSCD68DAPICASP1
CASP1
50μm
D
Relative expression of CASP1 in INS positive cells
Control
IL-1β Ab
TNFSF12 Ab
double Abs
SARS-CoV-2
E
CVB4
CVB4+IL-1β Ab
CVB4+TNFSF12 Ab
CVB4+double Abs
F
CVB4
G
HMGB1
ELANE
IL6ST
P=0.0563
BCL2L2
Normalized counts
Control
COVID
H
Differential signaling in human islets exposed to SARS-CoV-2
MIF-(CD74+CXCR4)
MIF-(CD74+CD44)
DLK1-NOTCH2
CD99-PILRA
APP-CD74
Mock
SARS-CoV-2
β to DC
β to immune progenitors
β to MΦ
β to T
Commun Prob
max
min
p-value
p>0.05
0.01<p<0.05
p<0.01
Differential signaling in human islets exposed to CVB4
NECTIN3-NECTIN2
NECTIN2-CD226
NAMPT-(ITGA5+ITGB1)
MIF-(CD74+CXCR4)
MIF(CD74+CXCR4)
MDK-SDC2
MDK-NCL
MDK-LRP1
MDK-(ITGA4+ITGB1)
HLA-E-KLRK1
HLA-E-KLRC1
HLA-E-CD8B
HLA-E-CD8A
HLA-C-CD8B
HLA-C-CD8A
HLA-B-CD8B
HLA-B-CD8A
HLA-A-CD8B
HLA-A-CD8A
GRN-SORT1
GDF15-TGFBR2
F11R-F11R
DLK1-NOTCH2
DLK1-NOTCH1
CDH1-CDH1
CDH1-(ITGAE+ITGB7)
CD99-PILRA
APP-CD74
Mock
CVB4

**Figure S6. TNFSF12 expression in human islets exposed to CVB4 or SARS-CoV-2, and VMI organoids with proinflammatory macrophages. Related to Figure 6.**

**(A)** Confocal images of TNFSF12 in human islets exposed to mock, CVB4 ($2x10^6$ PFU/ml) or SARS-CoV-2 (MOI=0.5). The inserts show a high magnification of cells. Scale bar= 25 µm.

**(B)** Confocal images of TNFSF12 in VMI organoids containing unstimulated or pro-inflammatory macrophages at day 7 after reaggregation. Scale bar= 20 µm.

**(C and D)** Confocal images (C) and quantification (D) of the CASP1 expression in $INS^+$ cells of VMI organoids exposed to SARS-CoV-2 (MOI=0.5) and treated with control, 10 µg/ml TNFSF12 blocking antibody, 5 µg/ml IL-1β blocking antibody or 10 µg/ml TNFSF12 + 5 µg/ml IL-1β blocking antibodies. The inserts show a high magnification of cells. Scale bar= 50 µm.

**(E and F)** Confocal images (E) and quantification (F) of the CASP1 expression in $INS^+$ cells of VMI organoids exposed to CVB4 ($2x10^6$ PFU/ml) and treated with control, 10 µg/ml TNFSF12 blocking antibody, 5 µg/ml IL-1β blocking antibody or 10 µg/ml TNFSF12 + 5 µg/ml IL-1β blocking antibodies. The inserts show a high magnification of cells. Scale bar= 50 µm.

**(G)** Normalized counts of pyroptosis associated genes expression in control or COVID-19 samples examined by GeoMx transcriptomic assays. Each dot represents one count in each ROI.

**(H)** Cell chat analysis showed the interactions from β cells to immune cell subpopulations, including DC cells, immune progenitors, T cells and macrophages in human islets exposed to SARS-CoV-2 (MOI=1) or CVB4 ($2x10^6$ PFU/ml).

N=3 independent biological replicates. Data was presented as mean ± STDEV. ****P* < 0.001.

**Table S1. Patient information. Related to Figures 1, 2, 6 and Figures S1, S2, S3, S6.**

**Table S2. Antibodies used for immunocytochemistry, intracellular flow cytometry analysis. Related to STAR Methods.**

| Usage | Antibody | Clone # | Host | Catalog # | Vendor | Dilution |
|---|---|---|---|---|---|---|
| Immunostaining | Polyclonal Guinea Pig Anti-Insulin | Polyclonal | Guinea Pig | #A0564 | Dako | 1:500 |
| Immunostaining | Glucagon Rabbit Ab | Polyclonal | Rabbit | #2760 | Cell Signaling | 1:1000 |
| Immunostaining | Polyclonal Rabbit Anti-Somatostatin | Polyclonal | Rabbit | #A0566 | Dako | 1:1000 |
| Immunostaining | Human CD31/PECAM-1 Antibody | Polyclonal | Sheep | #AF806 | R&D Systems | 1:1000 |
| Immunostaining | Purified anti-human CD68 Antibody | Monoclonal | Mouse | # 333802 | Biolegend | 1: 100 |
| Immunostaining | Cleaved Caspase-1 (Asp297) | Monoclonal | Rabbit | # 4199 | Cell Signaling | 1: 200 |
| Immunostaining | hPDX-1 Affinity purified goat igG | Polyclonal | Goat | # AF2419 | R&D Systems | 1:500 |
| Immunostaining | Nkx6.1 (D8O4R) Rabbit mAb | Monoclonal | Rabbit | #54551 | Cell Signaling | 1: 500 |
| Flow Cytometry | APC anti-mouse/human CD11b Antibody | Monoclonal | Rat | #101212 | Biolegend | 1: 50 |
| Flow Cytometry | APC anti-human CD206 (MMR) Antibody | Monoclonal | Mouse | #321109 | Biolegend | 1:50 |
| Flow Cytometry | APC anti-human CD14 | Monoclonal | Mouse | #301808 | Biolegend | 1:100 |

| | | | | | | |
|---|---|---|---|---|---|---|
| GeoMx | Insulin Monoclonal Antibody (ICBTACLS), Alexa Fluor™ 488 | Monoclonal | Mouse | # 53-9769-82 | Thermo Fisher Scientific | 1:200 |
| GeoMx | Cytokeratin, pan Antibody (AE-1/AE-3) [DyLight 594] | Monoclonal | Mouse | # NBP2-33200DL594 | Novus Biological | 1:200 |
| Immunostaining | caspase-1 Antibody (14F468) | Monoclonal | Mouse | #sc-56036 | Santa Cruz | 1:200 |
| Immunostaining | Enterovirus (Concentrate) | Monoclonal | Mouse | #M7064 | Dako | 1:500 |
| Immunostaining | CD163 (D6U1J) Rabbit mAb | Monoclonal | Rabbit | #93498 | Cell Signaling | 1:200 |
| Immunostaining | Anti-PRSS1 antibody produced in rabbit | Polyclonal | Rabbit | #HPA063471 | Sigma Aldrich | 1:500 |
| GeoMx | Purified anti-Cytokeratin 19 | Monoclonal | Mouse | #628502 | Biolegend | 1:1000 |
| Immunohistochemistry | Human B7-1/CD80 MAb (Clone 37711) | Monoclonal | Mouse | # MAB140-100 | RnD | 1:500 |
| Immunostaining | Alexa Fluor 488 AffiniPure Donkey Anti-Guinea Pig IgG (H+L) | Polyclonal | Donkey | #706-545-148 | Jackson Immunoresearch Labs | 1:500 |
| Immunostaining | Donkey anti-Mouse IgG (H+L) Highly Cross-Adsorbed Secondary Antibody, Alexa Fluor 594 | Polyclonal | Donkey | #A-21203 | Thermo Fisher Scientific | 1:500 |
| Immunostaining | Donkey anti-Rabbit IgG (H+L) Secondary Antibody, Alexa Fluor 594 conjugate | Polyclonal | Donkey | #A-21207 | Thermo Fisher Scientific | 1:500 |
| Immunostaining | Donkey anti-Rabbit IgG (H+L) | Polyclonal | Donkey | #A-31573 | Thermo Fisher | 1:500 |

| | | | | | | |
|---|---|---|---|---|---|---|
| | Secondary Antibody, Alexa Fluor 647 conjugate | | | | Scientific | |
| Immunostaining | Donkey anti-Mouse IgG (H+L) Secondary Antibody, Alexa Fluor 647 | Polyclonal | Donkey | #A-31571 | Thermo Fisher Scientific | 1:500 |
| Immunostaining | Donkey anti-Goat IgG (H+L) Cross-Adsorbed Secondary Antibody, Alexa Fluor 647 | Polyclonal | Donkey | #A-21447 | Thermo Fisher Scientific | 1:500 |
| Immunostaining | Donkey anti-Sheep IgG (H+L) Cross-Adsorbed Secondary Antibody, Alexa Fluor 647 | Polyclonal | Donkey | #A-21448 | Thermo Fisher Scientific | 1:500 |
| Immunostaining | Donkey anti-Mouse IgG (H+L) Highly Cross-Adsorbed Secondary Antibody, Alexa Fluor Plus 405 | Polyclonal | Donkey | #A48257 | Thermo Fisher Scientific | 1:500 |

**Table S3. Primers used for qRT-PCR. Related to STAR Methods.**

| **Primer name** | **Sequence** |
|---|---|
| *ACTB-Forward* | *CGTCACCAACTGGGACGACA* |
| *ACTB*-Reverse | *CTTCTCGCGGTTGGCCTTGG* |
| *ETV2-F* | *GAAGGAGCCAAATTAGGCTTCT* |
| *ETV2-R* | *GAGCTTGTACCTTTCCAGCAT* |
